# Interplay between cytokine and FGF2 signaling in induction of entosis and vasculogenic mimicry response in glioblastoma

**DOI:** 10.1101/2025.02.09.637275

**Authors:** Tae-Yun Kang, JinSeok Park, Alper Dincer, Janani Baskaran, Chunxiao Ren, Soraya Scuderi, Liang Yang, Declan McGuone, Kathryn Miller-Jensen, Jennifer Moliterno Günel, Murat Günel, Andre Levchenko

**Affiliations:** Systems Biology Institute, Yale University, West Haven, CT 06516, USA; Department of Biomedical Engineering, Yale University, New Haven, CT 06520, USA; Cancer and Blood Disease Institute, Children’s Hospital Los Angeles, CA 90027, USA; Keck School of Medicine, University of Southern California, Los Angeles, CA 90033, USA; Department of Neurosurgery, Yale School of Medicine, New Haven, CT 06520, USA; Department of Neurosurgery, Tufts Medical Center, Boston, MA 02111, USA; Program in Neurodevelopment and Regeneration, Yale University, New Haven, CT 06520, USA; Child Study Center, Yale University, New Haven, CT 06520, USA; Department of Pathology, Yale University, New Haven, CT 06520, USA

**Keywords:** Vasculogenic Mimicry, Glioblastoma, Entosis, Tumoroid, Macrophage, Cell-Cell Communication

## Abstract

Tumor vascularization is critical to survival of cancer cells, but is frequently perturbed leading to disorganized angiogenesis and emergence of alternative means of delivery of oxygen and nutrients, such as vasculogenic mimicry (VM). Understanding of VM and its relationship to endothelial vascularization has been hampered by the lack of comprehensive combination of *in vivo* clinical data and relevant *in vitro* models. We address this challenge by analyzing glioblastoma (GBM) tumors and clinically isolated cancer cells. This analysis strongly suggests a key role of macrophage-induced controlled cell death in emergence of VM. The results further point to entosis of cancer cells as a critical intermediate state in this process, enabled by mechano-chemical cell heterogeneity. We find evidence that macrophages can regulate endothelial angiogenesis and VM as two alternative vascularization mechanisms. These results reveal mechanistic underpinnings of VM and pave the way to predictive analysis of tumor progression.

## Introduction

Since Judah Folkman first proposed inhibiting tumor growth by targeting tumor-associated blood vessels^1^, anti-angiogenic therapy has rapidly evolved, uncovering the molecular mechanisms driving angiogenesis^2–5^. However, clinical trials reveal that these approached have disappointedly limited overall survival benefits ^4,6^. Tumors often resume growth after an initial response, indicating potential resistance to anti-angiogenic treatment. This resistance may arise from factors, such as heterogeneous response of endothelial cells (EC) or redundant pro-angiogenic pathways^7^. In particular, tumors can reduce dependence on angiogenesis by employing alternative vascularization strategies^8–11^ such as Vasculogenic Mimicry (VM).

VM is a fascinating and still-poorly understood process in which non-endothelial cells self-organize into vessel-like conduits that can enhance delivery of oxygen or nutrients into the tumor interior. VM was first identified in uveal melanoma^12^ and has since been associated with poor outcomes in various aggressive cancers^13–16^. Since VM uniquely occurs in an endothelial cell-free manner, it has been seen as potentially providing a key explanation for the limited efficacy of anti-angiogenesis drugs in cancer treatment.

The cellular origins and mechanisms of formation of VM structures are a subject of debate and active investigation. The initial and still frequently favored mechanism postulates emergence of VM structures from cancer stem cells (CSC). CSC or stem-like CD133+ cells, including a subset of vascular endothelial-cadherin (CD144)-expressing cells^17,18^, can directly differentiate into endothelial cells, subsequently forming vessel-like structures^19–21^. However, while some antiangiogenic therapies do target pathways identified in VM derived from CSCs^22–25^, in certain instances they have the opposite effect of enhancing rather than inhabiting VM formation^26,27^. These results have prompted the search for alternative mechanisms driving VM formation across various tumors, irrespective of tissue-specific genetic signatures of CSCs^28–30^. In particular, it has been proposed that VM structures can arise from self-organization of macrophages into inter-connected networks resembling vascular beds^31,32^. These diverse perspectives result in varied opinions on the definitive markers and methods for distinguishing VM^17,20,33,34^. Commonly, VM identification *in vivo* relies on detecting dilated lumens and thick glycoprotein, Periodic Acid Schiff (PAS)-positive, ECM, the markers that are noticeably distinct from capillary vessels. Depending on the mechanism, when CSCs or macrophages differentiate into EC-like cells, the lumen lacks the typical EC markers like CD31 or CD34, indicating incomplete EC differentiation.

In addition to the still outstanding question of the origin and mechanisms of VM, a key challenge still not addressed in investigation of this phenomenon is its relationship to angiogenesis leading to endothelial vascularization of the tumor mass. In particular, it is not clear if angiogenesis and VM occur independently of each other or they might represent alternative mechanisms of oxygen and nutrient supply, preferentially occurring in different local tissue micro-environments. Related to this, it is not well understood whether the triggers for angiogenesis and VM are distinct and potentially mutually exclusive.

The current lack of consensus about the origin of VM and its triggers point to the possibility that many alternative mechanisms can underly this complex phenomenon *in vivo*. However, they also may reflect the conceptual and technological focus on the traditional mechanisms of emergence of cellular structures, such as cell differentiation and self-organization. As a consequence, tools and approaches focused on such mechanisms are frequently transferred to the analysis of VM origins, ignoring the possibility of new mechanisms that may emerge in the context of tumor growth. Furthermore, the studies of VM rarely integrate the analyses on multiple scales, which bring together the molecular, cell and tissue levels of investigation with the examination of the related clinical samples and patient histories. Here, we employ a wide array of tools to undertake such a multi-scale analysis finding evidence of a new mechanism of VM formation in human glioblastoma (GBM).

In the analysis presented here, we use intra-operatively obtained live GBM cells and tissue samples, and an established cell line, to investigate the mechanisms of VM formation. To explore the mechanisms of VM, we used a wider range of methods, including modeling VM in GBM tumoroids supplemented with heterotypic cells and the analysis of clinical samples and images of GBM tumors. Using this diverse array of tools, we find evidence of a new mechanism of VM formation in GBM. Our results suggest that VM can form through entosis-mediated death of a subset of GBM cells, which can be triggered or enhanced by a signaling cross-talk between cancer cells and macrophages. Our analysis also indicates that angiogenesis-mediated endothelial vascularization and VM are not independent processes, but rather can serve as prominent alternative strategies enabling tumor survival and growth.

## Results

### Vasculogenic mimicry is induced in GBM tumors and GBM tumoroids in the 3D GBM modeling micro-device

We addressed tumor vascularization in GBM using both inter-operatively obtained live patient cancer cells and the clinical samples and data available for these patients. In this analysis we contrasted the behavior of cancer cell tumoroids grown in defined conditions and co-cultured with human macrophages and/or endothelial cells with the clinical information, imaging data and histological and immuno-histochemical analysis of tumor sections (Fig. 1A). This analysis explored both the natural occurrence of distinct vascularization outcomes in clinical samples and their emergence in the corresponding controlled 3D models. Furthermore, we analyzed in depth the putative molecular mechanisms leading to VM formation using a commercial GBM cell line, A172 (PTEN loss of function mutation, also present in the GBM cells from patient 580), and validated the resulting insights in clinical samples.

**Figure 1.**
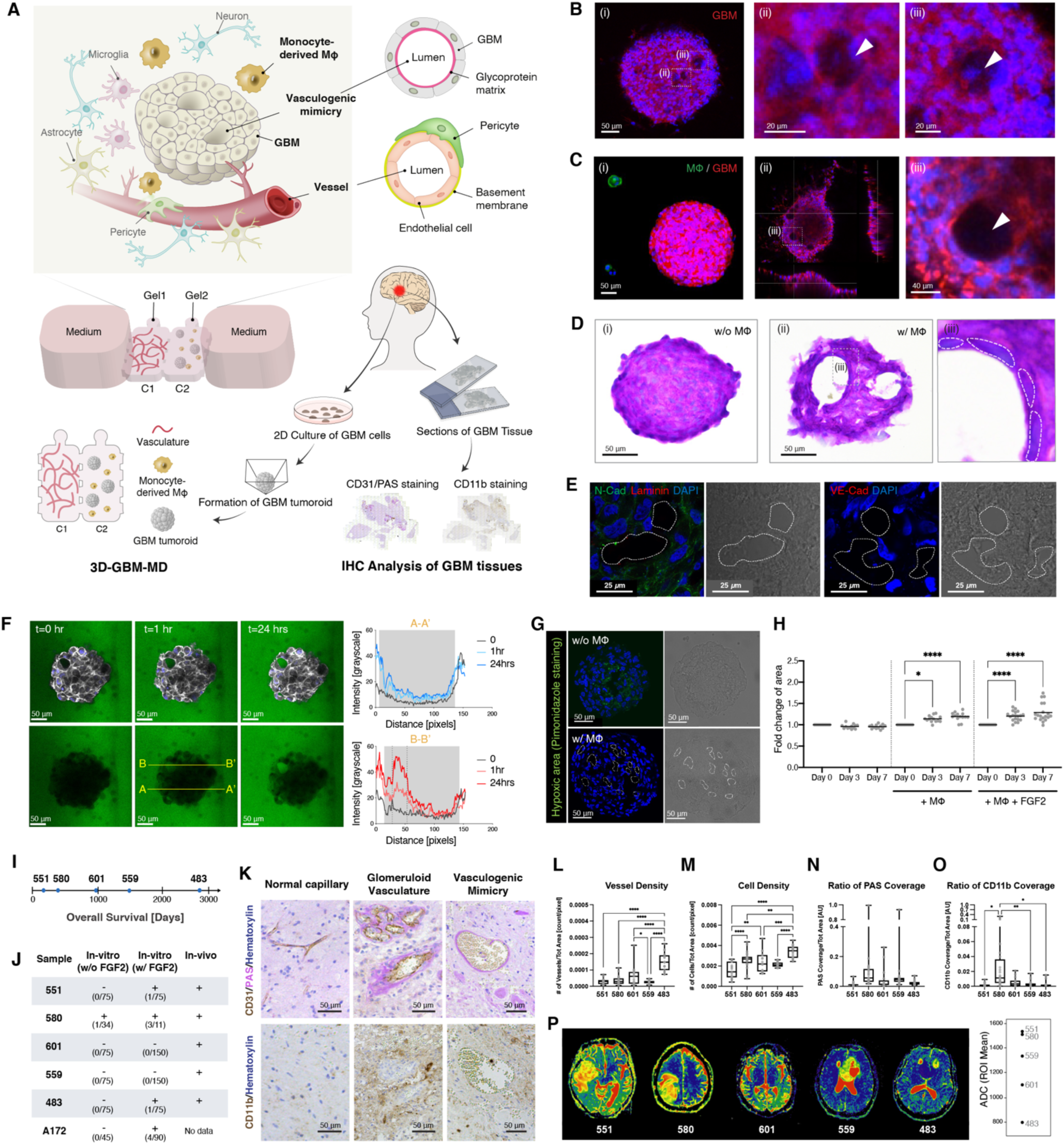
Vasculogenic Mimicry (VM) Formation in Multicellular *In vitro* and *In vivo* Environments. (A) Multiple approaches used in this study to investigate vasculogenic mimicry using cells and tissue sections derived from the same patient tumors, with the live cell experiments performed in GBM tumoroids grown in a highly controllable microfabricated platform (3D-GBM-MD) also accommodating models of human vasculature and macrophages. Human brain microvascular endothelial cells (HBMEC) were encapsulated within a fibrin gel in the chamber C1, while macrophages differentiated from U937 and glioblastoma tumoroids were encapsulated within a mixture of agarose and collagen gel in the chamber C2 of 3D-GBM-MD. (B) A representative image of a GBM tumoroid in 3D-GBM-MD (chamber C2 in (A)) after 5 days of culture in the absence of macrophages (i), with close-up views of cells hosted inside of other enlarged cells (ii &iii) indicated by white arrowheads; F-actins (red) and nuclei (blue) were stained with Alexa Fluor™ 647 Phalloidin and Hoechst 33342, respectively. (C) A representative image of a GBM tumoroid in 3D-GBM-MD after 3days culture in the presence of macrophages (i), with close-up views of cavity formations (ii &iii) indicated by white arrowheads; F-actins (red) and nuclei (blue) were stained as in (B). Macrophages were pre-labeled with CellTracker (green). (D) Hematoxylin (purple) and PAS (pink) staining of GBM tumoroid cross-sections after 3days culture without (i) and with macrophages (ii); PAS-coated lumen (iii) indicating formation of a vasculogenic mimicry vessel (indicated schematically in (A)). (E) Immunostaining of GBM tumoroid cross-sections after 5 days culture with macrophages: N-Cadherin (green), laminin (red), nucleus (blue) are shown on the left; VE-Cadherin and nucleus (blue) on the right along with phase contrast images; white dashed contours indicate cavities. (F) Representative FITC-dextran diffusion images (left) and intensity profiles (right) across ‘A-A’’and ‘B-B’ lines, passing through a cavity; Grey shades in the profiles represent tumoroid regions; dashed lines indicating start point and end point of the internal cavity; F-actins (white) and nuclei (blue) are indicated. (G) Representative pimonidazole staining (green) indicates the hypoxic cores in GBM tumoroids after 5 days culture without (top) and with (bottom) macrophages; white dashed contours indicate cavities. (H) Dynamic analysis of GBM tumoroid sizes from day 0 to day 7 under various culture conditions, with each tumoroid size individually normalized to its initial size on day 0. Data are presented by scatter dots representing individual tumoroids with the lines indicating the mean value. P-values were calculated using one-way ANOVA with Tukey’s multiple comparison test. (I) Overall survival (OS) of the five human patients who contributed cells and tissues to this study. (J) Status of VM induction using cells from individual patients or the A172 cell line after 5 days culture in the *in vitro* 3D-GBM-MD, along with VM observations in tissue samples from these patients. The status is marked with ‘+’ (presence) or ‘-’ (absence) and the specific number of positive cases out of the total number of tumoroids, as shown in parentheses (n/m). (K) Representative histological images from GBM patient tissues displaying three vessel types: capillary vessels glomeruloid vessels, and vasculogenic mimicry vessels (visualized and analyzed using CD31, PAS, and hematoxylin staining, top row). Distribution of monocytes or macrophages (CD11b and hematoxylin staining, bottom row, measured in the adjacent sections). (L) Vessel density, (M) cell density, (N) fraction of the PAS-positive area, and (O) fraction of CD11b coverage in tissues from human patients. Data are presented by box and whiskers with all individual points representing each image. P-values were calculated using one-way ANOVA with Tukey’s multiple comparison test. pixel to distance ratio: 1.075 pixels/*µ*m, pixel aspect ratio: 1 (P) The fMRI images of the patient tumors used in the analysis represented as the Apparent Diffusion Coefficient (ADC) maps (left) for each patient and quantified 3D volumetric ADC mean values (right). Tumoroids in (B) and (C) were made of primary GBM cells from human patient 580, while those in (D)-(H) were made of A172 cell line.

We performed tumoroid analysis in a new 3D GBM modeling micro-device (3D-GBM-MD) that permits close examination of tumoroid re-organization through confocal imaging and enables analysis of the effects of paracrine cell-cell communication (Figs. 1A, left & S1A). In different versions of this experimental setup, we examined the effects of co-culture with human macrophages and endothelial cells, as key non-cancer components of GBM tumors. Our initial focus was on the effect of macrophages. We found that without macrophages, tumoroids frequently contained enlarged cells encapsulating another cell (Figs. 1B & S1B top). However, with macrophages present, these cell-in-cell (CIC) structures were either absent or observed with much lower frequency. Instead, we observed emergence of hollow cavities within the GBM tumoroids (Figs. 1C &S1B bottom). Importantly, we found that the complementation of the medium with FGF2 (also known as bFGF, a growth factor implicated in GBM stemness^35^) strongly enhanced the formation of these vacuous structures. Intriguingly, these cavities exhibited the hallmarks of VM. In particular, histological staining of tumoroid sections (Fig. 1D) showed that the walls demarcating the cavities were positive for Periodic Acid-Schiff (PAS) and thus enhanced expression of glycogen-rich extracellular matrix (ECM), a key *in vivo* indicator of VM^15,36^. The lumens of the cavities were cell-free, and the cells on the abluminal side were positive for N-Cadherin, a junction protein typically expressed in GBM cells, yet they showed no detectable VE-Cadherin, an endothelial cell marker, or laminin, a component of the capillary basement membrane (Fig 1E). These results suggested that the cavities forming in tumoroids under the paracrine influence of macrophages have hallmarks of VM and that they were lined with GBM cells expressing VM markers.

We further examined whether the VM-like structures in tumoroids might have functional roles in enabling growth and survival of GBM cells. It is hypothesized that VM yields channels that can mediate replenishment of oxygen and nutrients at the interior of the tumor mass. First, we examined if molecular transport, as measured by penetration of FITC-Dextran (70 kDa) into the tumoroids, would benefit from formation of VM-like structures. We found that whereas there was very limited penetration of FITC-dextran into tumoroids that were cultured without macrophages and not forming VM-like structures (Fig. S1C), and also into the cavity-free areas of the tumoroids (Fig. 1F: A-A’). On the other hand, there was a strongly enhanced transport of FITC-dextran through the cavity network into the interiors of tumoroids (Fig. 1F: B-B’). Functionally, the enhanced transport through VM structures is expected to lead to relief of hypoxia that can develop at the tumor mass. We indeed observed significantly diminished hypoxic core formation within tumoroids (Fig. 1G) and an increase of tumoroid size (Fig. 1H & S1D) in tumoroids cultured in VM inducing environment. These results suggested that, functionally, the network of cavities in GBM tumoroids has the properties expected of VM. Overall, these results strongly suggested that VM can indeed be modeled in GBM tumoroids complemented with macrophages and FGF2.

Experimental analysis suggested that VM structure could form in tumoroids generated from most of the patient samples and from the A172 cell line (Figs, 1I, J, S1E&F). We then examined whether this finding would be mirrored by the analysis of VM in the tissue sections (Figs. 1K-O) obtained from the same patient tumors and further supported by the magnetic resonance imaging (MRI) images of these tumors (Figs. 1P, S1G&H). To explore this, GBM tissue sections from the five human patients were stained in two adjacent sections of each sample for PAS/CD31 (VM and endothelial markers) and CD11b (a marker for monocyte-derived cells, the majority of which are macrophages in GBM). We could identify three types of vessels by their morphologies and staining patterns (detailed in the Methods section): capillary vessels (CD31+/PAS-), glomeruloid vessels (CD31+/PAS+), and VM vessels (CD31-/PAS+) (Fig. 1K). Importantly, VM vessels were observed in tissue sections from all five human patients (Fig. 1J). Furthermore, VM was particularly prominent in the tumors enriched with macrophages (as revealed by CD11b staining). In particular, GBM 580 cells, exhibiting the most prominent VM formation *in vitro*, also showed elevated levels of PAS-positive ECM and macrophage/monocyte presence in the patient tissue (Fig. 1N and O). Interestingly, GBM 580 cells were the only ones that did not require complementation of the medium with FGF2 for VM formation in tumoroids. Importantly, we did not find evidence of macrophages forming the vessel walls, although they were frequently adjacent to the vessels. We further found a range of tumor cell density values across the tumors from different patients. This range of cell densities was contrasted with the MRI data examining the Apparent Diffusion Coefficients (ADC) maps, which reflected the cell density value in the entire tumor (Fig. 1P) obtained pre-operatively for each patient (Figs. S1G&H). We found that these MRI results demonstrated strong consistency with the cell densities quantified from tissue sections (Fig. 1M). This result supported the experimental assumption that the tissues selected for histological analysis adequately represented the entire tumor of each patient.

Our results suggested an important role for macrophages in VM formation, both in the patient tumors and in the tumoroid models *in vitro*. However, the mechanisms of this putative macrophage effect were not clear. Furthermore, it was not clear how FGF2 might enhance VM in the tumoroid model and whether the CIC structures observed in the absence of macrophages might be related to formation of VM structures. We thus set out to address these questions and to explore whether the corresponding biochemical processes might constitute different sides of the same VM regulation mechanism.

### Imbalance in FGF2 responsiveness leads to Cell-in-Cell formation, a prerequisite for VM formation

First, we explored whether CIC structure might be a precursor to VM and investigated the mechanisms of their formation. Prior studies suggested that CIC structures can emerge through a number of different mechanisms, all of which rely on the heterogeneity of cells in a population creating biochemical and, possibly, biomechanical asymmetry in adjacent cells^37,38^. For instance, in entosis, a stiffer cell can invade (and is engulfed by) a softer cell^39,40^. Such asymmetries may be amplified by external inputs. We hypothesized FGF2, in particular, can be such an input, with differential responsiveness of cells to FGF2 leading to enhanced biochemical differences between them and to enhanced incidence of CIC events (Fig. 2A). To test this hypothesis, we took advantage of chemotactic properties of FGF2^41–43^. Specifically, we constructed and used a microfluidic device (Fig. 2B) similar to those introduced by us previously^44,45^ to enrich the particularly chemotactic cell sub-population, which we hypothesized would also display an increased FGF2 sensitivity. This chemotactic enrichment of A172 cells was performed under normoxic and hypoxic conditions. The resulting populations termed unsorted (US, i.e., the original, population prior to enrichment), FGF2-chemotactic under normoxia (F) and FGF2-chemotactic under hypoxia (FH) were then used to explore the CIC properties and their relevance to VM.

**Figure 2.**
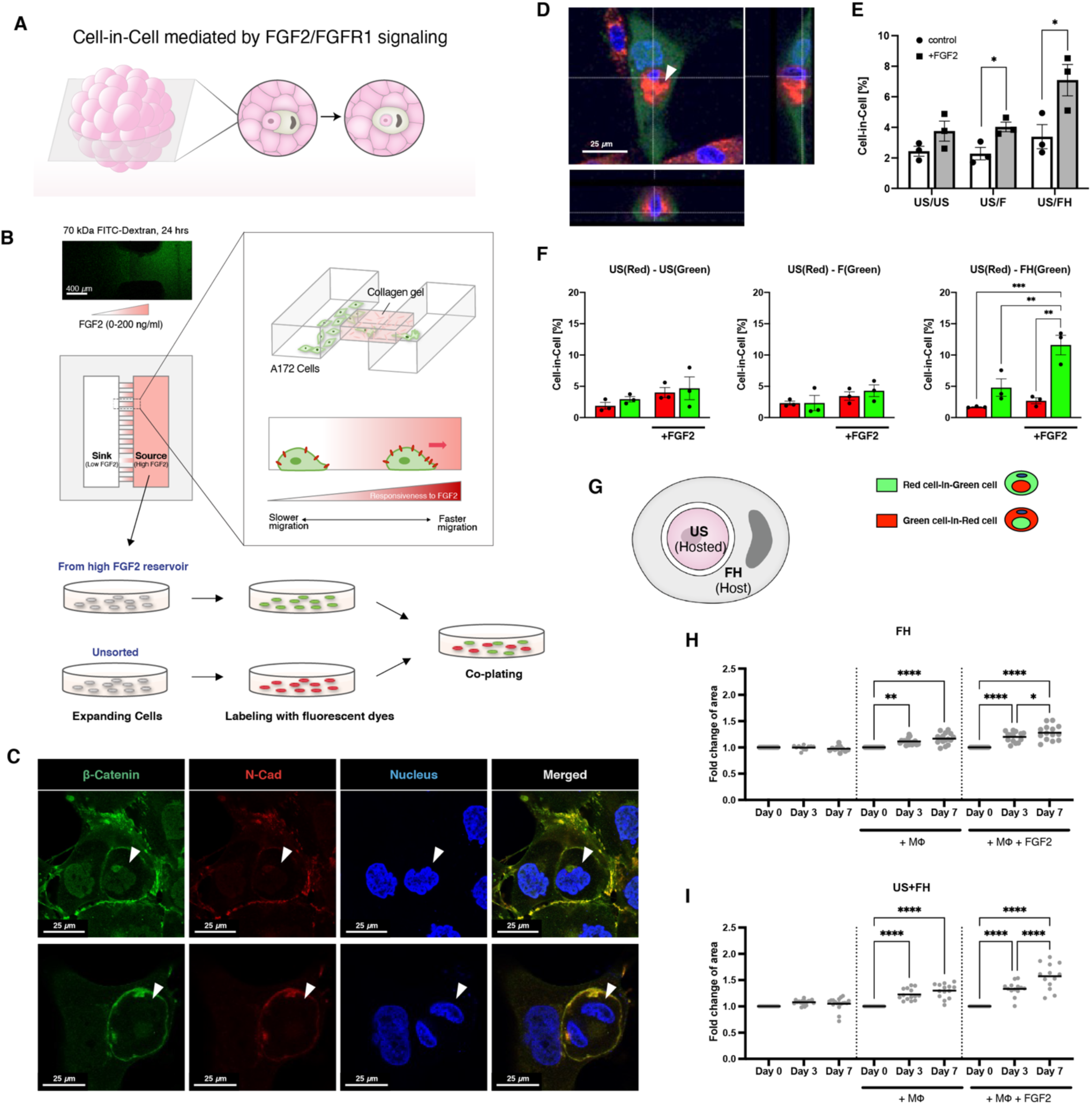
Cell-in-cell formation is enhanced by imbalance in FGF2 responsiveness. (A) A schematic summarizing the mechanism tested in experiments shown in this figure, illustrating formation of cell-in-cell (CIC) structures within 3D tumoroids promoted by FGF2/FGFR1 signaling. (B) A schematic of a microfluidic cell sorting chip designed for isolating cell sub-populations using chemotaxis up the FGF2 gradient yielding ‘sorted’ Fast (F) and Fast in hypoxic conditions (FH) cells, contrasted with ‘unsorted’ cells not subjected to chemotaxis-based separation. Also shown is the process of co-culture experiments using pre-labeled US cells with F or FH cells at 1:1 ratio. (C) Representative immunofluorescence images displaying CIC structures stained with beta-Catenin (green), N-Cadherin (red), and nuclear dye (blue), with examples of a quiescent cell within another cell (top row) and a cell undergoing mitosis inside another cell (bottom row) indicated by white arrowheads. (D) Confocal images showing a red-labeled cell completed hosted by a green-labeled cell in a 2D co-culture. (E) Percentage of CIC formations under various co-plating conditions: US/US, US/F, and US/FH, compared without (white bar) and with (grey bar) FGF2 treatment (n=3, mean ±SEM). Adjusted p-values were calculated using multiple paired t-tests with two-stage linear step-up procedure of Benjamini, Krieger and Yekutieli^46–48^. (F) Breakdown of CIC percentages from (E) based on host cell coloration (n=3, mean ±SEM). P-values were calculated using one-way ANOVA with Tukey’s multiple comparison test. (G) Experimental-derived schematic illustrating the US cell hosted by the FH cell. (H) Dynamic analysis of the sizes of GBM tumoroids made solely of FH cells, and (I) tumoroids of the US/FH mixture examined from day 0 to day 7 under various culture conditions, with each tumoroid size individually normalized to its initial size on day 0. For FGF2 treatment, 100 ng/ml was added into the medium. Data in (H) and (I) are presented by scatter dots representing individual tumoroids with the lines indicating the mean values. P-values were calculated using one-way ANOVA with Tukey’s multiple comparison test.

We found that CIC structures could be observed in both tumoroids and a 2D mono-culture of A172 GBM cells. The inner cells were encapsulated by the outer cells, with clearly defined interfaces marked ß-Catenin and N-Cadherin expression (Fig. 2C). The inner cells remained condensed and round (Fig. 2C, top row), and some even underwent mitosis within the outer cells (Fig. 2C, bottom row). We then used this analysis to explore the CIC formation by differentially labeling cell sub-populations. In particular, we co-cultured red-labeled US cells with green-labeled F or FH cells (Fig. 2B). Red-labeled US cells and green-labeled US cells was also co-plated as a control. 3D reconstructed confocal imaging was then used to identify and analyze CIC (Fig. 2D). The results revealed that FGF2 treatment increased the occurrence of CIC formation in all three co-culture combinations (Fig. 2E): US/US, US/F, and US/FH. However, the US/FH combination showed the highest rate of CIC formation, 3.4% without FGF2 and 7.1% with FGF2 treatment, while the rates of CIC formation in the US/F combination were similar to those in the US/US combination. Further analysis indicated that FH cells predominantly served as hosts (engulfing cells), while US cells were the hosted cells (Figs. 2F & G). The results held when the labels of these cells were switched (data not shown). Moreover, we observed a 3-fold increase in the rate of formation of tumoroids displaying VM when they formed from 1:1 mixture of US and FH cells vs. US cells alone. Finally, we found that organoids formed from these US/FH mixtures grew larger vs. US/US or FH/FH mixtures (Figs. 2H&I). These results suggested that cell heterogeneity rather than FGF2 sensitivity was the key parameter in both CIC and VM formation, and in VM function as an enabler of GBM cell survival and growth.

### Increased FGFR1 expression in FGF2-sensitive cells depends on receptor stabilization

We next explored the properties of FH cells, aiming to demonstrate more specifically their enhanced FGF2 sensitivity and the mechanisms driving their biochemical state. Immunoblotting revealed that FH cells expressed approximately 1.5-fold higher FGFR1 vs US or cells that did not migrate in the FGF2 gradient. In contrast, F cells showed no significant difference vs. US cells (Figs. 3A&B), suggesting that increased FGFR1 expression could be a key factor in the heightened sensitivity to FGF2, and supporting a key role of hypoxia in enhancing FGF2 sensitivity. Bulk RNA sequencing indicated that US, F, and FH cells were transcriptomically distinct populations, with F cells being more similar to US cells than to FH cells (Figs 3C-E). In agreement with our assumption about properties of FH cells, Gene Set Enrichment Analysis (GSEA) showed upregulation of genes related to FGFR1 and FGFR2 ligand binding and activation in FH cells compared to US cells (Figs. 3F, G, and S2A). FGF2 is known to bind FGFR1, FGFR2, and FGFR3, with FGFR1 having the highest specificity^49^. Bulk RNA sequencing data showed that FGFR2 is nearly absent, and FGFR1 is expressed approximately six times higher than FGFR3 in A172 cells (Fig. S2B). This result suggested that the responsiveness to FGF2 of A172 cells is primarily dependent on FGFR1. However, surprisingly, mRNA levels of FGF2 and FGFR1 were lower in FH cells compared to US cells (Figs. S2B & S2C).

**Figure 3.**
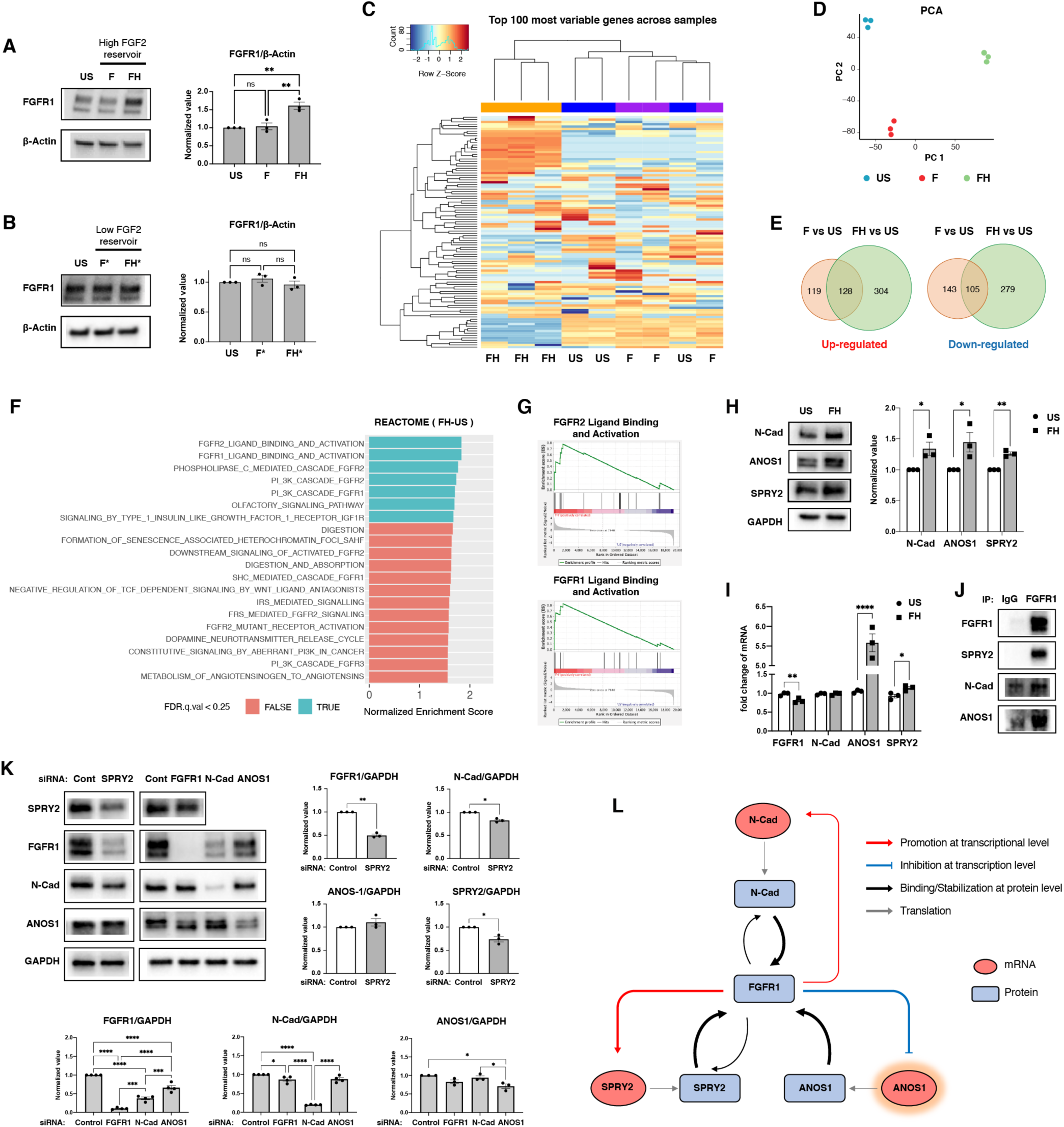
Higher FGFR1 level in sorted cells (FH) due to increased protein stability. (A) Representative immunoblots (left) of FGFR1 protein levels in unsorted A172 cells (US) compared to A172 cells from high FGF2 reservoirs sorted under normoxia (F) and hypoxia (FH), with quantification (right). β-Actin is used as a loading control (n=3, mean ±SEM). p-values were calculated using one-way ANOVA with Tukey’s multiple comparison test. (B) Representative immunoblots (left) of FGFR1 protein levels in unsorted A172 cells (US) compared to A172 cells from low bFGF reservoirs under normoxia (F*) and hypoxia (FH*), with quantification (right). β-Actin is used as a loading control (n=3, mean ±SEM). p-values were calculated using one-way ANOVA with Tukey’s multiple comparison test. (C,D): (C) Hierarchical clustering of the top 100 variable genes and (D) PCA plots from all gene expression data acquired via RNA-seq across US, F, and FH cell samples. n=3 biological replicates. (E) The Venn diagrams depicting the number of differentially expressed genes (DEGs; adjusted p-value < 0.05 and |log2(Fold Change)| > 1) in F and FH cells relative to US cells. (F) Reactome Gene Set Enrichment Analysis (GSEA) results in FH vs US cells. Positive normalized enrichment score (NES), enriched signature in FH. See also Figure S2A. (G) GSEA plots showing FGFR2 ligand binding and activation (top) and FGFR1 ligand binding activation (bottom) gene sets. (H) Representative immunoblots (left) of N-Cadherin, ANOS1, SPRY2 levels in US cells compared to FH cells, with quantification (right). GAPDH is used as a loading control (n=3, biological replicates, mean ±SEM). Adjusted p-values were calculated using unpaired multiple t-tests with two-stage linear step-up procedure of Benjamini, Krieger and Yekutieli. (I) Quantification of the fold change in FGFR1, N-Cadherin, ANOS1, and SPRY2 at the transcriptional level in US compared to FH, acquired from RNA-seq data (n=3 biological replicates, mean ±SEM). Adjusted p-values were calculated using unpaired multiple t-tests with two-stage linear step-up procedure of Benjamini, Krieger and Yekutieli. (J) Co-immunoprecipitation (Co-IP) assays of physical interactions of FGFR1 with SPRY2, N-Cadherin, and ANOS1. Negative control used IgG instead of FGFR1 antibody at a similar protein level. (K) Representative immunoblots (left) and quantification (right) demonstrating the effect of knocking down SPRY2, FGFR1, N-Cadherin, and ANOS1 via siRNAs on FGFR1 expression. GAPDH is used as a loading control (n=3 biological replicates, mean ±SEM). P-values were calculated using two-tailed paired t-test and one-way ANOVA with Tukey’s multiple comparison test to compare the differences by siSPRY2 and by multiple siRNAs, respectively. (L) A schematic overview of FGFR1 interactions with other proteins at both the transcriptional and protein levels.

Given that the increased FGFR1 protein expression in FH cells was not due to enhanced gene transcription, we investigated proteins known to physically interact with FGFR1—specifically, N-Cadherin and ANOS1, as well as SPRY2 known interact with the MET receptor —to determine if these interactions might stabilize the FGFR1 protein levels^50–52^. The expression levels of N-Cadherin, ANOS1, and SPRY2 were elevated in FH cells at the protein level, as shown in Figure 3H, paralleling the increase in FGFR1 (Fig. 3A). Notably, however, only the mRNA levels of ANOS1 and SPRY2 were higher in FH cells, as indicated in Figure 3I, with ANOS1 mRNA being approximately six times higher in FH vs US cells. Co-immunoprecipitation (Co-IP) experiments confirmed direct binding of N-Cadherin, ANOS1, and SPRY2 to FGFR1 (Fig. 3J). Among these proteins, SPRY2 showed the most pronounced binding to FGFR1. Knockdown of genes coding for these proteins led to a decrease in FGFR1 expression at the protein level (Fig. 3K). Intriguingly, FGF2 treatment resulted in decreased ANOS1 transcription (Fig. S2D), suggesting that the elevated ANOS1 mRNA level in FH cells is an inherent characteristic, rather than a consequence of enhanced FGF2-FGFR1 signaling. Overall, this analysis strongly suggested that FH cells exhibit higher FGFR1 protein levels not due to an increased FGFR1 gene transcription, but likely because of protein stabilization through direct interaction with N-Cadherin, ANOS1, and SPRY2 (Fig. 3L).

### FGF2-FGFR1 signaling regulates cell stiffness promoting entosis in heterogenous cell populations

We next investigated the mechanisms behind the propensity of FH cells with elevated FGF2-FGFR1 signaling to host US cells. We considered possible mechanism of entosis, a non-apoptotic cell engulfment process-involving the differential mechanical deformability driven by RhoA–ROCK signaling-mediated actomyosin contraction^38,40,53^, along with the clearance of apoptotic cells. This mechanism was favored since other potential mechanisms of cell engulfment by other cells were contradicted by experimental results. In particular, our observations ruled our phagocytosis of dying cells^54^ since that hosted cells were not commonly apoptotic (Fig. S3A) in the absence of macrophages. Our data also made it unlikely that entosis was promoted by mitotic cell division^55^ as, with few exceptions (Fig. 2C, bottom row), most hosted cells were non-mitotic (Fig. S3B). We also did not find evidence for dependence on reprogrammed glucose uptake^56^, leading to metabolic stress and AMP-activated protein kinase (AMPK) activation, since we did not find significant differences between US and FH cells in terms of phosphorylation of LDHA and AMPK after FGF2 treatment (Fig. S3C).

We next explored the possibility that the CIC formation could be due to direct regulation of RhoA– ROCK-MLC signaling-mediated actomyosin contraction through FGF2-FGFR1 signaling^57,58^. We and others have previously shown that MLC activity can be controlled by PI3K-Akt signaling and its activation of Rac1-mediated MLC phosphatase^59^, which can thus counteract ROCK-MLCK dependent MLC phosphorylation^60,61^. We thus explored activation of FGF2-induced signaling network in FH and US cells, finding that both Akt and the other canonical downstream kinase, Erk, were indeed activated in response to FGF2 in both these cell states. However, only Akt (but not Erk) displayed a significant increase in both baseline and FGF2 induced relative activation in FH vs. US cells. As expected, this led to a relative decrease in MLC2 phosphorylation (pMLC2) in FH vs. US cells, even though the total levels of MLC2 in FH cells displayed a relative increase vs US cells (Fig 4A). Of note, as seen in Figure S3D, the peak phosphorylation times after treatment varied, thus only the peaks following FGF2 exposure are displayed in Figure 4A: 15 minutes for pFGFR1, pAkt, and pErk, and 2 hours for pMLC2. Conversely, inhibition of FGFR1 kinase activity in FH cells resulted in down-regulation of Akt and Erk phosphorylation and up-regulation of pMLC2 (Fig. 4B). To provide further information on the relative roles of Akt and Erk in pMLC2 regulation, we pharmacologically inhibited each of the kinases (Fig. 4C), finding a reciprocal activity pattern (i.e., inhibition of Akt led to an increase in pErk and vice versa). However, only inhibition of Akt led to up-regulation of pMLC2 regardless of the pErk status, consistent with the negative role of this pathway in MLC2 activation.

**Figure 4.**
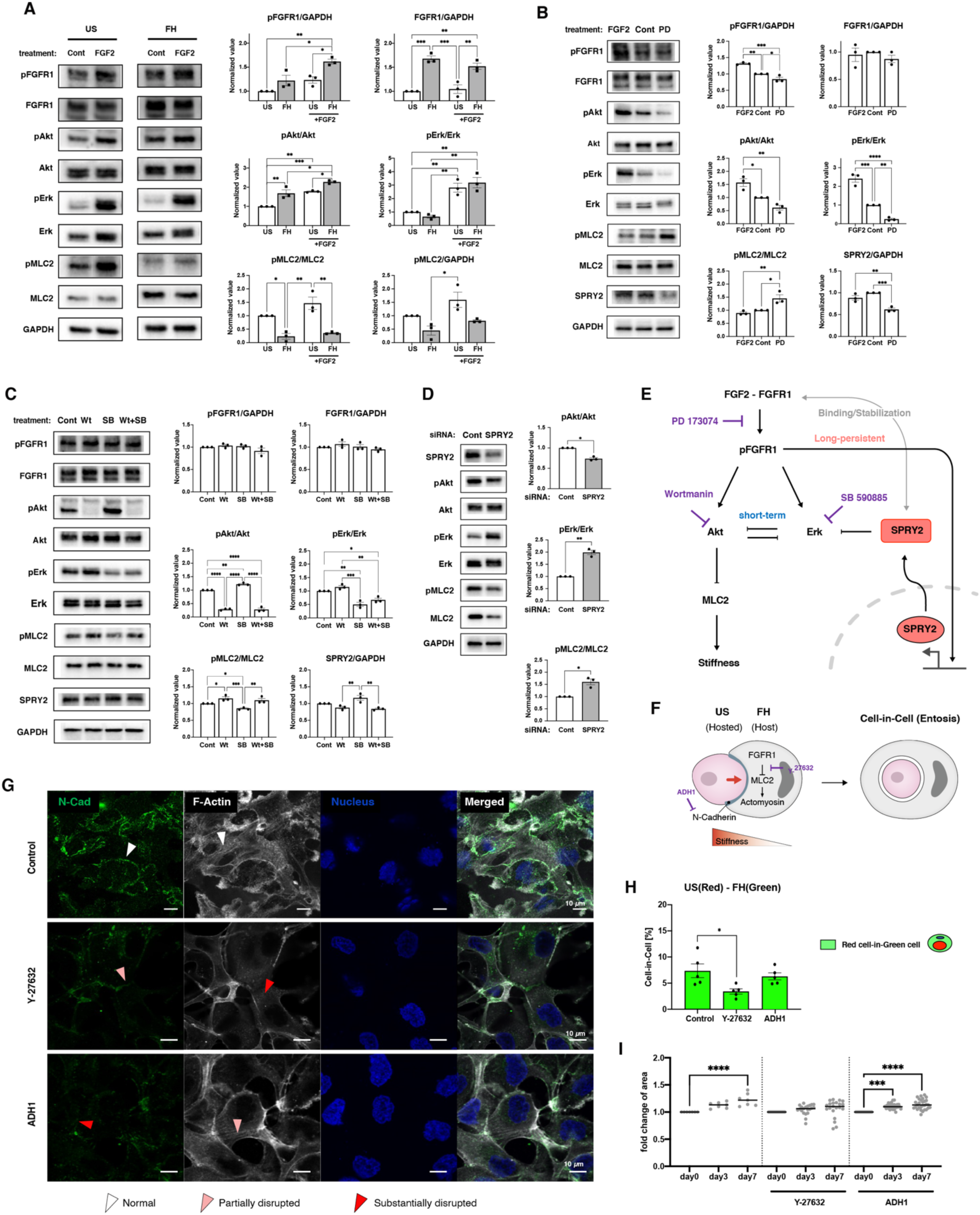
The signaling network regulating entosis and CIC formation in response to FGF2. (A) Representative immunoblots showing activation of signaling pathways in US and FH cells in response to FGF2 (100 ng/ml) treatment (left), with quantification (right). GAPDH is used as a loading control (n=3 biological replicates, mean ±SEM). Only peak values of the assayed phosphor-proteins after FGF2 treatment are displayed and analyzed: 15min for pFGFR1, pAkt, and pErk, and 2hr for pMLC. See also Figure S3D for detailed temporal dynamics. P-values were calculated using two-way ANOVA with Tukey’s multiple comparison test. (B) Representative immunoblots (left) showing the signaling outputs without and with sustained inhibition of autophosphorylation of FGFR1 24 hours using PD173074 (0.1 µM) treatment in US cells (left), with quantification (right). The conditions and analysis details are otherwise as in (A). GAPDH is used as a loading control (n=3 replicates, mean ±SEM). P-values were calculated using one-way ANOVA with Tukey’s multiple comparison test. (C) Representative immunoblots showing sustained inhibition of Akt phosphorylation with wortmannin (2 µM) and of Erk phosphorylation with SB 590885 (1 µM), 24 hours post-treatment, both individually and in combination in US cells (left), with quantification (right). The conditions and analysis details are otherwise as in (A). GAPDH is used as a loading control (n=3, mean ±SEM). P-values were calculated using one-way ANOVA with Tukey’s multiple comparison test. (D) Representative immunoblots (left) showing the effect of knocking down of SPRY2 via siRNAs on downstream pathways of FGF2-FGFR1 signaling, with quantification (right). GAPDH is used as a loading control (n=3, mean ±SEM). p-values were calculated using two-tailed paired t-tests. (E) Overview of the signaling mechanisms controlling cell stiffness via FGF2-FGFR1 signaling. (F) A schematic illustrating the (CIC) formation caused by a stiffness imbalance between US and FH, regulated by FGF2-FGFR1 signaling. (G) Immunostaining images showing actin bundles and N-Cadherin treated with Y-27632 (0.5 µM) and ADH1(20 µM) in A172 cells. (H) Percentage of CIC formations observed in A172 cells treated with Y-27632 (0.5 µM) and ADH1 (20 µM). Red-labeled US and green-labeled FH cells were co-plated at a 1:1 ratio (n=5, mean ±SEM). The target of each drug in entosis regulation is indicated in (F). P-values were calculated using one-way ANOVA with Tukey’s multiple comparison test. (I) Dynamic analysis GBM tumoroids in the US/FH mixture, co-cultured with macrophages in 3D-GBM-MD from day 0 to day 7 in the presence of Y-27632 (0.5 µM) and ADH1 (20 µM). Each tumoroid size was individually normalized to its initial size on day 0. Data are presented by scatter dots representing individual tumoroids with mean lines. p-values were calculated using one-way ANOVA with Tukey’s multiple comparison test.

Of interest, we also found that the level of SPRY2 that we previously found to be controlled by FGFR1 on the genetic level (Figs. 3I&S2D) was up-regulated in response to Erk (but not Akt) inhibition (Fig. 4C). This is consistent with prior reports suggesting that SPRY2 can form a negative feedback with Erk, serving to limit the kinase activity^62–65^. To further explore this possibility, we examined signaling activity in SPRY2-knockdown FH cells (Fig. 4D). We found that pErk was indeed up-regulated whereas pAkt was reciprocally down-regulated and pMLC2 was up-regulated, again consistent with the negative effect of Akt activation on MLC2 phosphorylation. Overall, our results suggested that SPRY2 has a dual role in FGF2 signaling in FH cells. Its expression both can stabilize FGFR1 at the protein level and limit activation of Erk but not Akt, resulting in higher basal and induced Akt phosphorylation and lower MLC phosphorylation in FH vs. US cells (Fig. 4E), without an appreciable difference on pErk. Lower MLC phosphorylation in turn results in lower relative cell stiffness in FH cells and in entosis (Fig. 4F), if FH cells are co-localized with cells with lower FGFR1 expression and FGF2 sensitivity (present in US population).

These results suggested that enrichment of softer cells with decreased pMLC2 can enhance entosis-mediated CIC formation and promote VM. A decrease rather than increase in biomechanical cell asymmetry can be achieved by inhibition of pMLC2 across the cell population. We therefore tested the effect of suppression of pMLC2 by inhibition of the upstream kinase, ROCK (Fig. 4F). We found that this perturbation partially disrupted N-cadherin-mediated cell-cell contacts (Fig. 4G) and significantly inhibited CIC (Fig. 4H). VM and tumoroid growth in the presence of macrophages were also strongly inhibited (Fig. 4I). This effect was not due to the inhibition of cell contacts as direct N-cadherin inhibition did not alter these functional outcomes, suggesting the key influence of cell mechanics rather than adhesion. Furthermore, partial inhibition of ROCK has been reported to increase rather than decrease cell proliferation in diverse cell types, including GBM^66–68^. Therefore, the effect of ROCK inhibition on tumoroid growth was not due to inhibition of cell cycle, but rather decreased VM formation. Taken together, these results suggest that bio-mechanical cell asymmetry is indeed involved in entosis-mediated CIC phenomenon, leading to VM.

### Paracrine cell communication with GBM cells can promote M1-like macrophage polarization influencing GBM phenotypes

Our results still left open the questions of how CIC phenomenon can lead to formation of VM and how macrophages might mediate this process. To address these questions on a mechanistic level, we performed single-cell RNA sequencing (scRNA-seq) analyzing a 1:1 US:FH cell mixture in 3 conditions: 1) 2D culture, 2) 3D tumoroid culture without macrophages (-MΦ), and 3) 3D tumoroid culture with macrophages (+MΦ) (Fig. 5A). Distinct culture conditions led to widely separated clustering in the latent gene expression space, as shown in Figures 5B and S4A. Labeling US cells with tdTomato-N1 and then tracking the expression of this gene led to its detection in approximately 10% of US cells in scRNA-seq data, reflecting the dropout in single cell experiments. Nevertheless, the pattens of gene expression for all the highly expressed genes in tdTomato-N1-positive and -negative cells were essentially identical reflecting the same expression patterns in the US cells with detectable tdTomato-N1 and US with nondetectable tdTomato-N1:FH cell mixture and thus the predominance of culture conditions over the cell state in defining gene expression this experiment (Figs. S4B-E).

**Figure 5.**
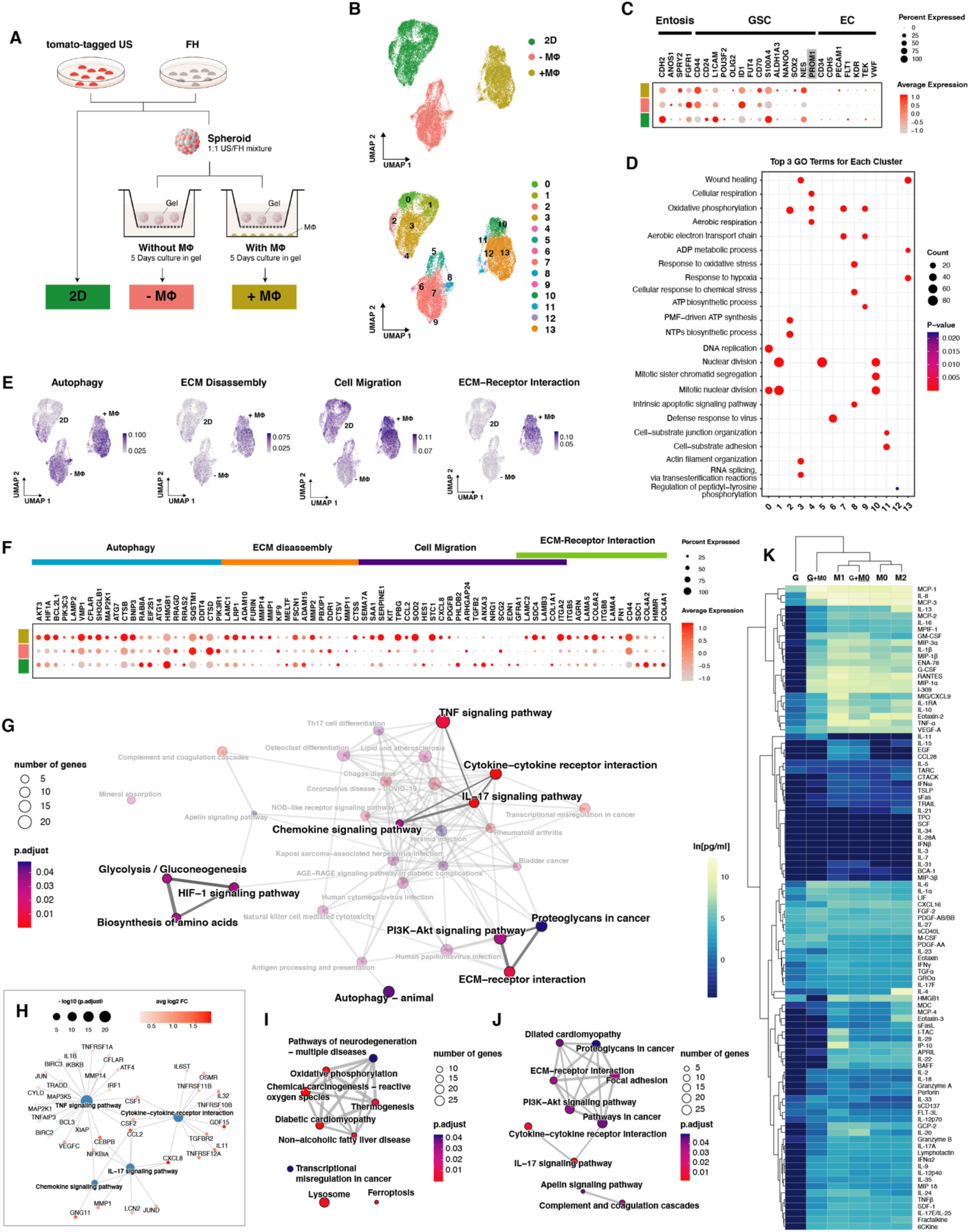
Comprehensive analysis of GBM transcriptional changes and functional associations in a macrophage-conditioned environment. (A) Experimental design to analyze GBM transcriptomic profiles induced in diverse culture conditions for 5 days. GBM tumoroids composed of US and FH (1:1 ratio) were collected from the gel and dissociated into single cells for analysis following incubation in the -MΦ and +MΦ conditions. (B) UMAP visualization of single-cell transcriptomes, color-coded by culture conditions (top) and by clusters (bottom) (C) Dot plot displaying entosis, glioblastoma stem cell (GSC) and endothelial cell-specific gene expression signatures across culture conditions. The dot size represents the percentage of cells in which the gene of interest was detected, with the color indicating the gene’s scaled average expression in those cells. PROM1 is shaded in gray to indicate that the gene was not detected. (D) Top 3 Gene Ontology (GO) terms (BP: Biological Process) enriched in each cluster. See also Figure S4G for GO terms from comparisons between conditions. Cutoff : p<0.05; adjusted p (p.adj) < 0.05. (E) Module scores displayed on UMAP for feature expression of marker genes of selected GO and GSEA terms: Autophagy, ECM disassembly, Cell Migration, ECM-Receptor interactions. (F) Dot plot displaying selected GO and GSEA terms-specific gene expression signatures across culture conditions. The dot size represents the percentage of cells in which the gene of interest was detected and color indicates the gene’s scaled average expression in those cells. (G) GSEA enrichment map displaying gene sets grouped into a network with edges connecting overlapping gene sets, from the KEGG database of DEGs found to be upregulated in +MΦ tumoroid condition compared to 2D culture. The dot size represents the number of genes in the term of interest and color indicates the p.adj-value. Cutoff: p<0.05. (H) Functional association networks between signature genes specific to TNF signaling pathway, cytokine-cytokine receptor interaction, IL-17 signaling pathway, and chemokine signaling pathway selected from (G). The size of blue dots represents log10(p.adj-value) of the term of interest and the shade of the red dots represents average log2(Fold Change) of each gene. (I) GSEA enrichment map from the KEGG database of DEGs upregulated in -MΦ tumoroids condition compared to 2D culture (the dot size and color are as in (G)). (J) GSEA enrichment map from the KEGG database of DEGs upregulated in +MΦ compared to -MΦ tumoroids conditions (the dot size and color are as in (G)). (K) Hierarchical clustering of 96-plex secretion profiles from GBM cells and macrophages under various culture conditions: G: GBM cells only; **<u>G</u>**+M0: GBM cells co-cultured for 3 days with M0 macrophages; M0: newly differentiated macrophages; M1: macrophages polarized by IFN-γ and LPS; M2: macrophages polarized by IL-4 and IL-13; G+**<u>M0</u>**: M0 co-cultured with GBM cells for 3 days. DMEM was used as the basal medium for all conditions. Color shades indicate the concentration values according to the scale. See the full result in Table S2.

Analysis of the expression of individual genes (Figs. 5C &S4D) suggested an increase in the level of SPRY2 in a subset of tumoroid cells in the presence vs absence of macrophages, with both conditions resulting in higher expression levels vs 2D culture. On the other hand, ANOS1 and FGFR1 expression levels were maximized in tumoroids in the absence of macrophages, which however may not reflect the FGFR1 protein levels that can be stabilized by SPRY2 and ANOS1, as discussed above. The expression patterns also suggested an increase in stemness markers^69–71^ in tumoroids in the presence of macrophages, particularly SOX2 and CD44, and their upstream regulator, CD70^72^. The mesenchymal stem cell marker CD44 was detected in almost all cells, whereas the expression of a more general stemness maker SOX2 was more limited. This suggested that mesenchymal differentiation of stem and mature GBM cells was enhanced by macrophages. Importantly, this effect of macrophages is consistent with their M1 polarization, known to promote the mesenchymal fate of GBM cells^73^, something that we tested in experiments described below. Finally, we detected no markers of endothelial differentiation, which again argued against VM structures arising through endothelial-like state of cancer cells.

Further analysis showed that cell populations in each condition can from distinct clusters in the UMAP latent space, with the detailed lists of the top five genes from each culture condition and from each cluster provided in Figures S4E and S4F, respectively. This dataset and its analysis for enrichment of GO terms (Figs. 5D & S4G) further supported mesenchymal differentiation of GBM cells in the presence of macrophages. In particular, this was suggested by the enrichment for GO terms related to cell adhesion and wound healing and the scores related to ECM remodeling and migration in +MΦ condition (Figs. 5E&F), both of which are hallmark features of cells with the mesenchymal phenotype. The GSEA analysis (Figs. 5G-J) further suggested a change in the metabolic state associated with hypoxia in tumoroid vs 2D culture, which could be accounted by both the actual increase in the hypoxic conditions in 3D and possibly the effect of macrophages on stabilization of HIF-1α via the NF-κB-dependent pathway^74^. Indeed, there was a strong enrichment of cytokine signaling, and more specifically TNFα and IL-17 signaling pathways in the culture conditions that included macrophages (Figs. 5G&H). This was consistent with putative secretion of both cytokines by macrophages, during which IL-17 can both induce and synergize with TNFα induction of multiple downstream pathways^75,76^. This result further suggested that macrophages were polarized towards the M1 fate in the presence of GBM cells. The data further suggested that proteoglycan, ECM-reception interaction, PI3K signaling GO terms are up-regulated in the presence of macrophages (Figs. 5G &J).

Support for macrophages polarization to M1 was further provided by the results of a multiplex secretion assay for 96 cytokines based on color-coded polystyrene beads (Fig. 5K). The analysis of secretion spectra of naive (M0), M1 and M2-polarized macrophages in isolated cultures, co-cultured macrophages and GBM tumoroids and GBM tumoroid controls revealed that the secretion pattern of macrophages co-cultured with GBM cells was clustered more closely with that of M1-like macrophages, which were induced by IFN-γ and LPS vs. M2-like macrophages, which were induced by IL-4 and IL-13. This was particularly evident in the secretion of similar levels of TNFα by M1 macrophages vs the M0 macrophages co-cultured with GBM tumoroids. The multiplex assay also revealed a change in the secretion of GBM cells co-cultured with macrophages vs. macrophage-devoid control. In particular, we found a strong up-regulation of secretion of IL-8 and CCL2 (MCP-1) (Figs. 5K & S5A). These results corroborate the prior report^77^ that GBM cells can release IL-8 and CCL2, which promote secretion of TNFα by macrophages and are in agreement with other reports suggesting paracrine interactions between GBM and macrophages^73,78^.

ELISA assays confirmed the cytokine secretion patterns observed in the multiplex analysis. In particular, we found that TNFα secretion was about six-fold higher in M1-like macrophages (M1) and macrophages co-cultured with GBM cells (G+**<u>M0</u>** _D3), compared to the baseline (M0). This peak occurred on day 3 (G+**<u>M0</u>** _D3) and decreased by day 5 (G+**<u>M0</u>**_D5) (Fig. S5B). Similar peaks and subsequent declines by day 5 were observed in the secretion patterns of other cytokines from the multiplex assay (Fig. S5A), with some exceptions. For instance, TRAIL secretion (Fig. S5C) was minimal and undetectable by both the multiplex and ELISA assays. These secretion patterns were not affected by the specifics of the culture medium (Figs. S5D&E), suggesting the key influence of cell-cell communication.

Overall, our studies suggest that GBM and macrophages can engage in paracrine signaling leading to polarization of M0 macrophages towards the M1 fate. This polarization, through secretion of TNFα and possibly other cytokines can promote mesenchymal transition of GBM and VM formation. We next set out to explore the putative mechanisms connecting the pro-inflammatory signaling to CIC and VM phenomena.

### Lysosomal membrane permeabilization-like leakage of entotic vacuole leads to joint cell death of host and hosted entotic cells

Our results suggested that macrophages can promote conversion of CIC structures to vacuous cavities that can ultimately form VM-like structures, in a macrophages (MΦ) dependent fashion. One way this process might occur is through elimination of CIC cells by regulated cell death, through a deleterious effect of macrophages (Fig. 6A). Indeed, we found that co-incubation with macrophages resulted in an increased extent of apoptosis of both the host and hosted cells in CIC structures, with close to 100% of hosted cells dying (Fig. 6B), consistent with the switch from CIC to VM structures that we observed in prior experiments (Figs. 1B&C).

**Figure 6.**
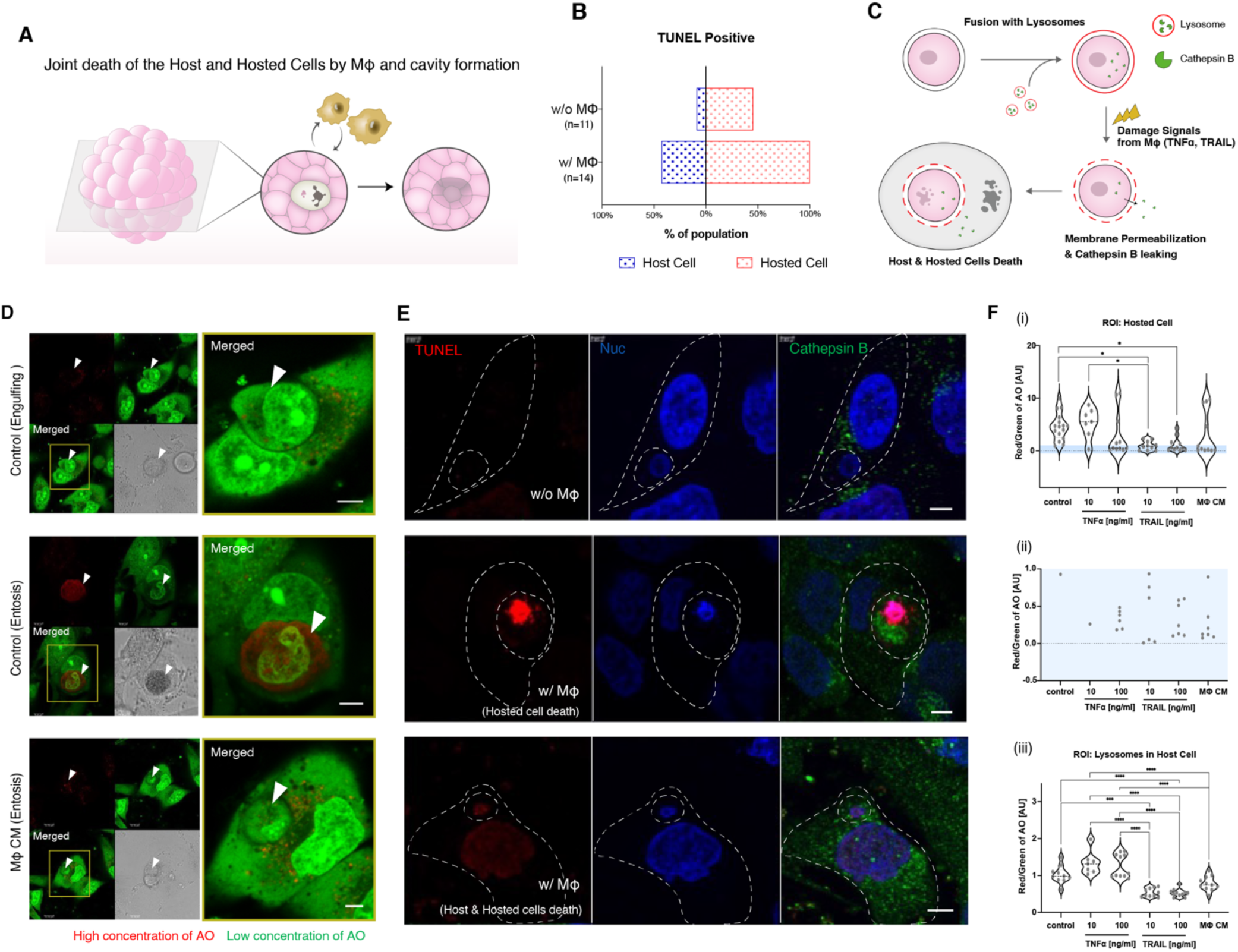
Cavity formation in tumoroids resulting from the joint cell death of both the host and the hosted cells due to lysosomal membrane permeabilization-induced leakage of the entotic vacuole. (A) The schematic summary of experimental observations, illustrating the cavity formation from the joint cell death of the host and hosted cells within 3D tumoroids in the presence of macrophages. (B) Percentages of TUNEL-positive host and hosted A172 cells in CIC structures, in the absence and presence of macrophages. (C) Schematic representation of the proposed selective clearance of CIC structures: joint cell death due to lysosomal membrane permeabilization-induced leakage of the entotic vacuole, trigged by damage signals from macrophages. (D) Images of Acridine Orange (AO) staining with close-up views (yellow squares) on the right: red in acidic membrane-bound organelles such as lysosomes and entotic vacuoles; green in cytosol and in leaking membrane-bound organelles. MΦ CM: conditioned medium from macrophages co-cultured with GBM. Scale bar: 5 µm (E) Representative images of co-staining of TUNEL and Cathepsin B localization in A172 CIC structures, in the presence and absence of macrophages. Scale bar: 5 µm (F) Ratios of Red/Green fluorescence from AO staining, serving as an indicator of membrane integrity for entotic vacuoles around hosted cells in CIC structures (i & ii) and lysosomes within the host cells (iii), following exposure to damage signals: TNFα, TRAIL, and MΦ CM. Image (ii) is a magnified view of the blue-shaded area in (i). Data are presented by violin plots with all points representing ROI of individual CIC structures. P-values were calculated using one-way ANOVA with Tukey’s multiple comparison test.

To further explore the possible mechanisms of macrophages-induced cell death, we focused on our finding that macrophages co-incubated with GBM cells can secrete TNFα and other potentially cytotoxic cytokines (Fig.5K). Cell death induced by TNFα is dependent on permeabilization of lysosomes releasing Cathepsin B (Cat B) in the cytosol^79,80^. We hypothesized that cells may be sensitized to this process if the permeabilized lysosomes contain hosted cells in CIC (Fig. 6C). To experimentally test this hypothesis, we conducted a series of tests. The use of Acridine Orange (AO), a metachromatic lysosomotropic dye, revealed lysosomal accumulation of AO molecules in the entotic vacuoles of host cells fully enclosing the hosted cells (Fig. 6D, middle row). This accumulation was not observed in cases where cell engulfment was still in progress (Fig. 6D, top row). When macrophage-conditioned medium (MΦ CM), harvested from GBM-macrophage co-cultures, was added to the cell culture, it induced permeabilization of entotic vacuoles. This led to a predominant green fluorescence signal within the entotic vacuoles, resulting from the dispersion of Acridine Orange (AO) (Fig. 6D, bottom row). In the absence of macrophages, the Cat B signal was weak (Fig. 6E, top row). In cases where only the hosted cell was dying in the presence of macrophages, Cat B was primarily localized within the hosted cells (Fig. 6E, middle row). Conversely, when both the host and hosted cells were TUNEL-positive, indicating apoptosis, Cat B was present in the cytoplasm of both cells (Fig. 6E, bottom row). It is noteworthy that in our scRNA-seq data (Fig. 5F), the Cat B gene (CTSB) was upregulated in the presence of macrophages (+MΦ). In combination, these data supported putative involvement of lysosomal permeabilization in cell death of cells in CIC states.

Among the cytokines that could contribute to creating an inflammatory environment, TNFα and TRAIL are known to induce lysosomal membrane permeabilization (LMP) through the downstream pathways of death receptors^79^. To ascertain the effects of TNFα and TRAIL, and to assess their impact relative to damage signals secreted by macrophages, we treated cells with cytokines and measured the red to green fluorescence intensity ratios (R/G) in entotic vacuoles surrounding hosted cells and in lysosomes within host cells, using AO staining. These measurements were taken for cells treated with TNFα (10ng/ml, 100ng/ml), TRAIL (10ng/ml, 100ng/ml), and MΦ CM. Our findings demonstrated that TNFα, TRAIL, and MΦ CM all reduced the R/G values in entotic vacuoles, indicating their increased permeabilization (Fig. 6F(i)). While the effect of TNFα varied, a high concentration (100 ng/ml) brought the median R/G value close to zero, similar to the effect of MΦ CM treatment. TRAIL was able to effectively lower the R/G ratio even at the lower concentration (10 ng/ml) (Fig. 6F(ii)). A similar distribution pattern of R/G values, quantified from lysosomes in host cells (Fig. 6F(iii)), suggested that entotic vacuole permeabilization mechanisms are similar to the established effects of cytokines on LMP.

Although both TNFα and TRAIL promoted death of CIC cells, only TNFα was secreted by macrophages co-cultured with GBM cells (Figs. 5K, S5A-E). Therefore, our experiments suggest that TNFα is the key factor inducing lysosomal membrane permeabilization (LMP)-like leakage in entotic vacuoles within CIC structures, leading to the initiation of VM.

### Glioblastoma exploits inflammatory environments in shifting its vasculature from normal capillaries to pathological vessels

Our results strongly suggest that macrophages can promote VM. However, these cells can also control morphogenesis of endothelial vascular networks^81^. We thus next utilized the other capability of the 3D-GBM-MD device: analysis of endothelial network formation in the presence of microenvironment defined by A172 GBM tumoroids, macrophages and controlled extracellular milieu (Fig. 7A). As expected, we found that both VEGF addition alone and GBM tumoroid co-culture without endogenously added VEGF could promote vasculogenesis, yielding lumenized vascular networks There was an additive increase in endothelial vessel formation in response to a combined VEGF/tumoroid exposure. Strikingly, vasculogenesis promoted by tumoroids was strongly inhibited by co-culture with macrophages, with this effect enhanced when VEGF was added to the culture medium in the device. This inhibitory effect was consistent with our prior observation that macrophages co-cultured with GBM tumoroids can promote M1-like fate and enhance TNFα secretion (Fig. 5K), and the prior results from our and other groups suggesting that high TNFα doses can interfere with VEGF signaling in endothelial cells^81,82^. These results suggested that macrophages can have a dual effect, at least in the 3D-GBM-MD setting, of both promoting the VM-like phenotype and inhibiting endothelial vascularization. We then explored if this duality may translate into the *in vivo* GBM setting.

**Figure 7.**
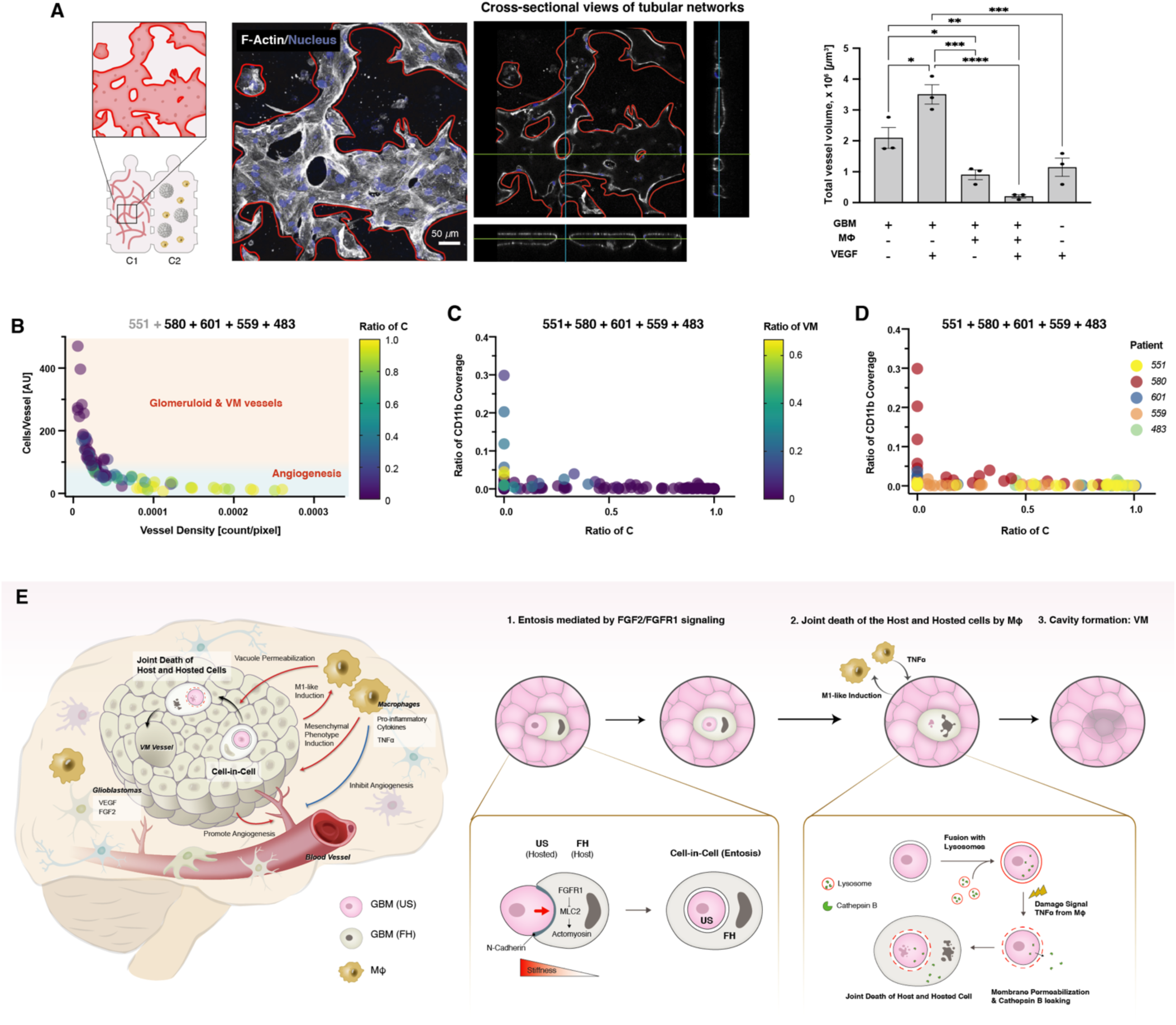
Shifting balance of vasculature type in GBM tissues from normal capillaries to pathological vessels under inflammatory environments. (A) Representative confocal images (left) showing endothelial cell-driven vasculogenesis in C1 chamber of 3D-GBM-MD platform (see Figure 1A) and quantified total vessel numbers (right) from 3 days culture in various conditions with GBM tumoroids and macrophages in the C2 chamber, further stimulated by VEGF treatment (n=3 biological replicates, mean ± SEM). P-values were calculated using one-way ANOVA with Tukey’s multiple comparison test. (B) Scatterplot relating the vessel density and the number of cells supported by each vessel in each analyzed location within a tumor tissue section. The color of each data point indicates the ratio of capillaries relative to all vasculature types in GBM tissues, including glomeruloid and vasculogenic mimicry vessels. Data from 87 locations across patient 580, 601, 559, and 483 are integrated into the plot. See also Figures S6E & F for plots that includes data from patient 551. See Figures S6G & H for individual plots for tissues obtained from each patient. (C) Scatterplot relating the ratios of capillaries and the ratios of CD11b coverages in each analyzed location within a tumor section. The color of each data point indicates the ratio of vasculogenic mimicry vessels relative to all vasculature types in GBM tissues. Data from all patients are integrated into the plot. See Figure S6I for individual plots from each patient. (D) Scatterplot of data shown in (C), indicating the individual tumors. (E) Graphical summary illustrating the interaction between glioblastoma cells, macrophages, and vasculature, leading to the emergence of vasculogenic mimicry. The proposed mechanism for vasculogenic mimicry formation involves entosis and lysosomal membrane permeabilization-like leakage from the entotic vacuoles within the tumor microenvironment triggered by the stress signals from the adjacent macrophages and enabled by differential mechanical properties of tumor cells that can depend on heterogenous growth factor signaling.

To establish the relative abundance of different types of vessels within patient samples, we re-scored the GBM tissue sections and quantified the numbers of capillary, glomeruloid and VM vessels in multiple locations (Figs. S6A-I). We found that the tumor from the patient 551 had a particularly low abundance of vessels of any type, which notably was associated with relatively much higher velocity and persistence of GBM cell migration (Fig. S6J) and lower cell density in the tumor (Fig. 1M), likely suggesting invasive cell spread and relatively low reliance on vascular cell support for this tumor. For the rest of the patients, there was a similar average abundance of glomeruloid vessels, but much more variable average abundances of both capillary and VM vessels. To ascertain the functionality of the vessels we explored the relationship between the vessel number and the number of tumor cells in the same area of the section, which are supported by these vessels. For 87 locations, for samples analyzed from all patients other than patient 551, we found a strikingly consistent inverse relationship between the vessel density and the number of tumor cells per vessel for each location (Fig. 7B). Importantly, we found the inverse relationship between vascular and tumor cell densities held for different types of vessels, not just capillaries. Furthermore, the capillary-rich regions predominantly constituted the high vascular density part of this relationship, whereas VM vessels tended to have lower density, with glomeruloid vessels falling in-between. The consistency of this dependency suggested that, functionally, all vessels constitute a continuum oxygen/nutrient delivery system, with VM vessels capable of nourishing larger number of cells per vessel, with lower vessel density vs. capillaries, likely due to their greater cross-section size. Indeed, we found that there was much greater average vessel size for glomeruloid and VM vessels vs capillaries (Fig. S6D).

We then tested whether the reciprocal dependency of angiogenic/vaculogenic vascularization vs. VM on macrophages, suggested by 3D-GBM-MD device experiments (up-regulation of VM and downregulation of capillary vessels), would hold *in vivo*. We again focused on the analysis of 108 locations in tumor samples, exploring the relationship between the capillary and VM formation and CD11b marker expression (Figs. 7C&D). We indeed found a universal inverse relationship between capillary vessel ratio and macrophage coverage for all the locations examined. On the other hand, the ratio of VM vessels increased with CD11b. Overall, these results suggested that GBM associated macrophages can promote VM formation and inhibit endothelial capillary vasculature formation, leading to a switch in the local mechanisms of oxygen and nutrient delivery.

## Discussion

VM continues to be an incompletely understood phenomenon that may be critical for cancer progression. Our study revealed a new mechanism of VM emergence supported by *in vitro* and *in vivo* results in human GBM. In contrast to the previous theories of how VM may develop, linking its onset to differentiation of cancer stem cells into endothelial-like cells and re-organization of newly differentiated or mature cells into vascular-like structures, we find evidence for the critical involvement of cell death in this process. More specifically, we find two key steps that may lead to the emergence of cell-free areas constituting lumens of VM vessels: 1) entosis driven by cell stiffness imbalance regulated in part by FGF2-FGFR1 signaling, and 2) joint cell death of host and hosted cells undergoing entosis, caused by LMP-like leakage from the entotic vacuole, triggered by TNFα secretion from M1-like macrophages (Figure 7E). The macrophage polarization is regulated by their proximity to and interaction with tumor cells. This mechanism was suggested by an array of experiments with human GBM cell lines, including those freshly derived from patients, and consistent with the clinical data and samples obtained from these patients.

Beyond the mechanism of VM formation suggested by our results, our data helps address the question of whether VM and angiogenesis are independent processes that might be both activated by the lack of nutrients or oxygen in the tumor mass. Our data suggest that these processes can indeed co-occur in the same tumors, but the relative frequency of their occurrence may be tumor specific. Some tumors, including that from patient 483 had predominantly endothelial vascularization, while others, such that in patient 580 had a large fraction of VM vessels. Nevertheless, we also found that, across 87 distinct locations in tumor tissue sections of all patient tumors we tested, there was a universal relationship between the number of tumor cells per vessel (of any kind) and the vessel density (Fig. 7B). This relationship encompassed a smooth transition from an almost exclusively endothelial vasculature for high vessel density to almost exclusively VM at low vessel density. This result suggested that VM or glomeruloid vessels, which are larger in diameter, can support more cancer cells, including cells that are located further away from a VM vessel vs. the average distance from an endothelial capillary. This result lends further support to the functionality of VM in GBM. More importantly, we found another universal relationship that held true for 108 different locations across the patient samples: enrichment of VM in proportion to the local density of macrophages. This result strongly suggested an important role of macrophages as potential triggers of the switch from angiogenic vascularization to VM. This relationship was consistent with our *in vitro* 3D-GBM-MD tumoroid data suggesting that macrophages co-incubated with GBM cells can polarize into an M1-like state and, as a result, promote VM and inhibit endothelial vasculogenesis.

It is of interest to contrast the results of our study with prior reports on possible roles of tumor associated macrophages in VM. Our results across multiple assays and from the analysis of patient tumor samples support a crucial role played by these cells in formation of VM structures. They are consistent with prior suggestions that macrophages may induce VM onset^83^ but are in contrast (at least in GBM) to the suggestion that macrophage cells themselves can organize into VM-like tubular structures^31^. In particular, we did not find any evidence of macrophages forming vascular walls in tumor sections. Our results further suggest that macrophages can be induced by GBM cells to adopt the M1 polarization state, which in turn leads to secretion of TNFα and promotion of VM. Of interest, prior reports suggested that hypoxic conditions in tumor microenvironments can induce transition to M2 polarization, which is thought to be pro-angiogenic^84,85^. These findings again underscore that endothelial vascularization and VM could be alternative outcomes that may be dependent on specific local microenvironments and macrophage polarization status. GBM tumors are particularly enriched in macrophages that show a mixture of polarization states^73^, which may in turn explain co-existence of VM and endothelia in the same tumors.

Both angiogenesis and VM can serve as adaptive strategies enabling cancer cell survival in the context of hypoxia and lack of nutrients in growing tumors. Another strategy, particularly relevant to GBM, is local invasive spread and dissemination of cancer cells from the site of the primary tumor thus enabling cells to relocate to areas better suited for their survival. Our limited set of tumors revealed all these three strategies in different patient tumors. In particular, the cell density in tissue slices and the ADC values from MRI images from the whole tumors both suggested low cell density and low vascularization in the tumor of patient 551. On the other hand, the same data sources suggested high endothelial vascularization and high cell density in the tumor from the patient 483, and the presence of endothelial and VM structures and intermediate cell density in other tumors. Interestingly, these diverse tumor architectures are reflected in different behaviors of cancer cells isolated from the tumor mass cultured in biomimetic devices. Cell behavior in these devices is highly similar to that in brain slices and predictive of clinical outcomes^86^. We found that the cells from patient 551 displayed high persistence and speed of cell migration which may correlate with the invasive dissemination/escape strategy suggested by the analysis of clinical samples, the MRI imaging studies and the shortest survival of this patient. On the other hand, the cells from the almost exclusively angiogenic tumor (patient 483) had high migration speed but the lowest migration persistence, suggesting inefficient invasive escape capability, correlating with the longest survival. Finally, the other tumors that displayed effective mixed VM and endothelial vasculature, also yielded tumor cells that had both relatively lower speed and persistence characteristics.

Our work underscores the importance of GBM plasticity and heterogeneity^87^. In particular, we find that differences in the activation of the signaling networks, such as that triggered by FGF2, can result in a spectrum of mechanical cell properties controlled by pMLC. This variability within cancer cell population is essential for the successful entosis, implicated here in the initiation of VM. Notably, FGF2 is stabilized by hypoxia^88^. Therefore, as in the microfluidic device used here, there may be chemotactic recruitment of FGF2-sensitive and thus softer GBM cells to the areas of high FGF2 concentrations, which coincide with the hypoxic zones. This, in turn, can lead to co-localization of cells with different stiffness properties and the resulting entosis and VM in these areas.

How does the entotic cell death lead to formation of a vessel, apparently requiring coordinated death of multiple adjacent cells and stabilization of the resulting cavities within dense tissues? We favor the mechanism that has been previously invoked to explain waves of cell death through ferroptosis observed in development^89^. Indeed, sensitivity to ferroptosis is elevated in GBM due to overexpression of ferroptosis promoting genes in tumor cells^90,91^. Ferroptosis may be enhanced due to the high levels of accumulated glutamate, initially exported by GBM cells, which can be re-imported through the glutamate-cysteine exchanger, leading to cysteine depletion in some cells^92^. Importantly, ferroptosis can be also promoted by local hypoxia due to the positive effect HIF-1α in GBM cells^93^. Indeed, we found that expression of the key ferroptosis promoting genes was up-regulated in the 3D GBM culture vs. 2D, indicating possible effects of oxygen depletion and glutamate accumulation (Fig. 5I). Thus, entosis may be a trigger in the process that generates a wave of cell death in adjacent cells, which may lead to progressive expansion of a vessel-like structure, particularly in hypoxic areas. This is distinct from commonly postulated self-organization of pre-existing cells into vascular structures (as in vasculogenesis) or branched growth of new ones (as in angiogenesis). Proteoglycans, which can be secreted by GBM cells^94^ and macrophages^95,96^, can modulate FGF-mediated signaling^94,97^, consistent with upregulation of the GO terms for PI3K signaling along with proteoglycan in Figures 5G and 5J of this study. Importantly, proteoglycans may be crucial for stabilizing the ECM structure^98,99^, and are the key marker of the luminal VM surfaces. Therefore, the cavities forming by macrophage-mediated death of entotic GBM cells might be stabilized by increased proteoglycan secretion.

Overall, our study uncovers a new mechanism of VM formation in GBM that crucially depends on heterotypic cancer cell-macrophage interaction and associated regulated cell death. This mechanism is in contrast to all previous models of VM postulating processes critically dependent on cell differentiation and self-organization. The close integration of diverse data sets and, in particular, the linkage to the analysis of clinical samples and data can pave the way for better understanding of VM emergence and its clinical relevance.

## Limitations of the study

The work presented here aims at bridging the *in vitro* and *in vivo* characterization of VM and, in particular, the analysis in 3D of VM-like structures using markers previously employed *in vivo*^14,15,100^. However, our work does not fully unravel the progression of initial VM structures into more complex tubular networks or their integration with existing blood vessels. We hypothesize that this process can involve active recruitment of macrophages to the initial VM formation sites leading to a positive feedback driving vessel formation. Testing this hypothesis necessitates further detailed investigation into the nascent and developmental stages of VM. While we favor the mechanism in which waves of cell death through ferroptosis contribute to the progression of vasculogenic mimicry following the initiation by the death of cells in CIC states, further studies are needed, along with efforts to address technical challenges in monitoring this phenomenon in three dimensions. In addition, further analysis will reveal the role of deposited extracellular matrix in protecting the developing channels from collapse following the initial death mediated cavity formation. Another limitation of our study is the use of U937 cells as a uniform proxy for macrophages *in vitro*. This approach doesn’t reflect the diverse behaviors and characteristics of macrophages, which could significantly influence the outcomes, particularly in the context of patient-specific degree of correlation between CD11b coverage and the emergence of pathological vessels. Development of methods to track interactions between primary patient-derived GBM and macrophage cells will help uncover more a detailed picture of VM emergence and influence on individual patient responses.

## Resource Availability

### Lead contact

Further information and requests for resources and reagents should be directed to and will be fulfilled by the lead contact, Andre Levchenko.

### Materials availability

This study did not generate new unique reagents.

## Acknowledgements

We thank Joseph O’Brien from the Department of Neurosurgery at Yale School of Medicine for maintenance of the database.

We acknowledge the following grant support: Mia Neri Foundation (T.-Y.K.); Financial support of the Department of Neurosurgery, Yale University (to T.-Y.K., M.G., and A.L.); NCI U54 U54CA209992 (to A.L.)

## Author contributions

T.-Y.K., M.G., and A.L. conceived the study. T.-Y.K., J.P., and A.L. designed and optimized the experiments. T.-Y.K. performed and analyzed 2D and 3D in-vitro experiments, including the 3D GBM modeling micro-device, cell-in-cell formation, immunoblotting, immunostaining (ICC and IHC), real-time PCR, 96-plex secretion, and bulk/sc-RNA sequencing. A.D. analyzed MRI images. J.B. and K.M.-J. designed the ELISA assay, and J.B. performed and analyzed the ELISA assay. J.P. and C.R. performed and analyzed the cell migration assay. S.S. prepared cryosections of 3D tumoroids. L.Y. produced the tdTomato-N1 expressing A172 cell line. D.M. arranged the preparation of patient GBM cryosections and contributed to discussions on the IHC analysis method. J.M.G. and M.G. provided patient GBM tissues, GBM cells and clinical information. T.-Y.K. performed and analyzed IHC with patient GBM tissues. T.- Y.K. and A.L. wrote the manuscript, with all authors providing edits and comments.

## Supplementary Information document

**Document S1: Methods**

**Document S2: Figures S1-S6**

Figure S1 related to Main Figure 1

Figure S2 related to Main Figure 3

Figure S3 related to Main Figure 4

Figure S4 related to Main Figure 5

Figure S5 related to Main Figure 5

Figure S6 related to Main Figure 7

