## Supplemental Figures for "Interplay between cytokine and FGF2 signaling in induction of entosis and vasculogenic mimicry response in glioblastoma"

#### Figure S1 related to Main Figure 1

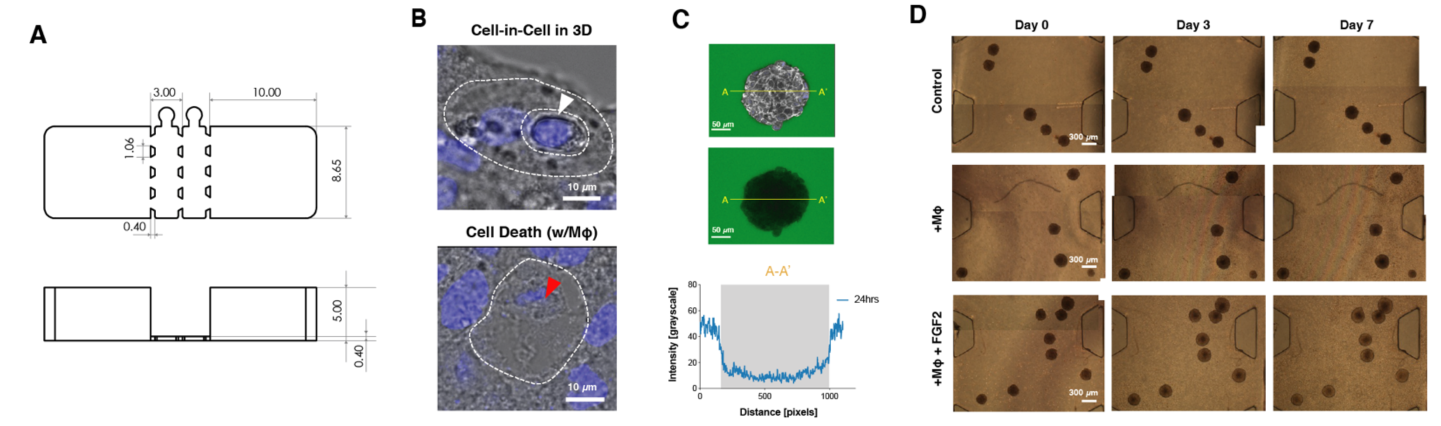


E

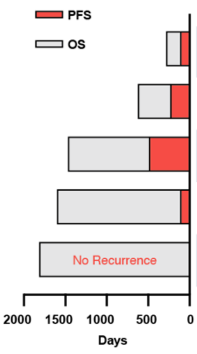


| Sample | Resection | Radiation (RT) | Temozolomide (concomitant with RT) | Temozolomide (adjuvant) | Bevacizumab | Immune-Checkpoint Inhibitor | Other | MGMT Methylation | IDH1 Mutation |
|---|---|---|---|---|---|---|---|---|---|
| 551* | Subtotal | + | + | - | + | - | - | + | - |
| 580* | Subtotal | + | - | - | + | + (Nivolumab) | - | - | - |
| 601 | Near total (97%) | + | + | + | + | - | Cediranib, Olaparib | - | - |
| 559 | Gross total | + | - (due to elevated LFTs) | + | - | - | - | + | + |
| 483 | Subtotal | + | + | + | + | - | - | + | - |

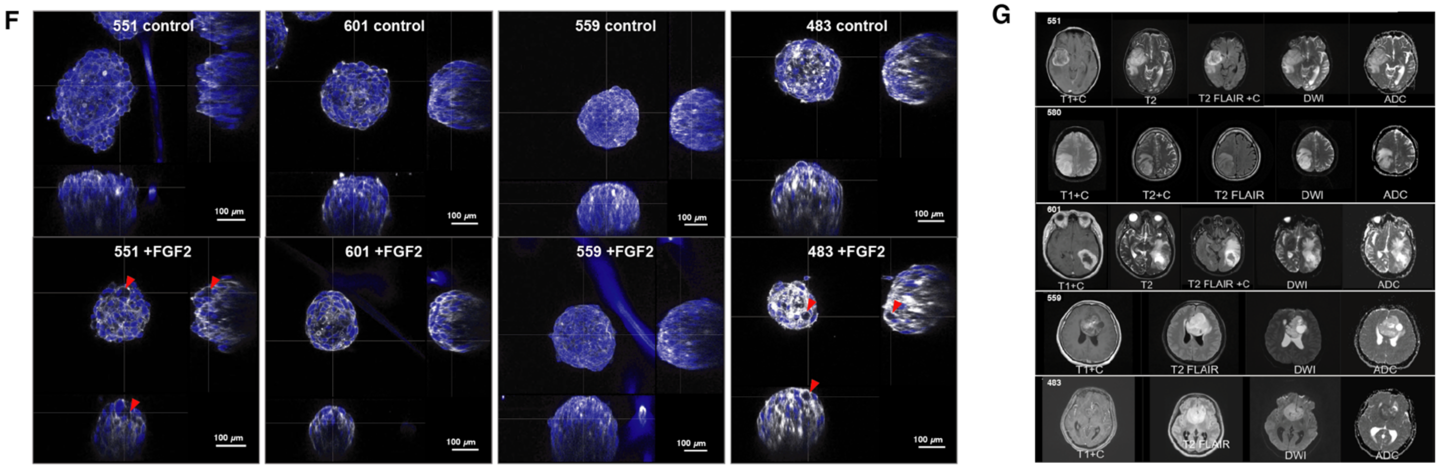


H

| Tumor ID | Location | Contrast | T2/FLAIR | DWI/ADC | ADC Max | ADC Min | ADC Mean |
|---|---|---|---|---|---|---|---|
| 551 | temporal | Ring-enhacing | Diffuse peritumoral edema | No diffusion restriction | 2784 | 740 | 1534 |
| 580 | parietal | Homogenous, Diffuse | Diffuse peritumoral edema | No diffusion restriction | 2219 | 576 | 1509 |
| 601 | parietal | Ring-enhancing | Diffuse peritumoral edema | No diffusion restriction | 1853 | 585 | 1101 |
| 559 | frontal | Scattered, Heterogenous | Minimal peritumoral edema, primarily intratumoral edema | No diffusion restriction | 2616 | 531 | 1333 |
| 483 | frontal | Minimal, Focused intratumoral necrotic cores | Minimal peritumoral edema, primarily intratumoral edema | Diffusion-restricting | 2574 | 365 | 796 |

**Figure S1. Details of GBM tumoroid culture in 3D TME-mimicking chip.**

(A) Dimensions of 3D-GBM-MD. Unit: mm

(B) Transmitted light images of cell-in-cell and clearance of a cell-in-cell structure in the presence of macrophages in GBM tumoroids captured by a confocal microscope.

(C) FITC-dextran diffusion images (top) and intensity profile (bottom) across A-A'; Grey shades in the profile representing tumoroid regions; F-actins (white) and nuclei (blue) were stained.

(D) Representative phase contrast images showing A172 tumoroids growth from day 0 to day 7 in C2 of 3D-GBM-MD under various conditions.

(E) Treatment summary and prognosis of each patient after surgical resection of tumors. OS, Overall Survival; PFS, Progression Free Survival. An asterisk (*) denotes patients for whom overall survival (OS) has been completed due to death.

(F) Three orthogonal views of 3D reconstructed confocal images displaying tumoroids made of primary GBM cells, observed both without and with FGF2 treatment in the presence of macrophages. Red arrowheads indicate cavity formation within the tumoroids.

(G) Preoperative Magnetic Resonance Imaging (MRI) characteristics of patients who contributed cells and tissues to this study.

(H) Interpretation of the MRI images of (G).

### Figure S2 related to Main Figure 3

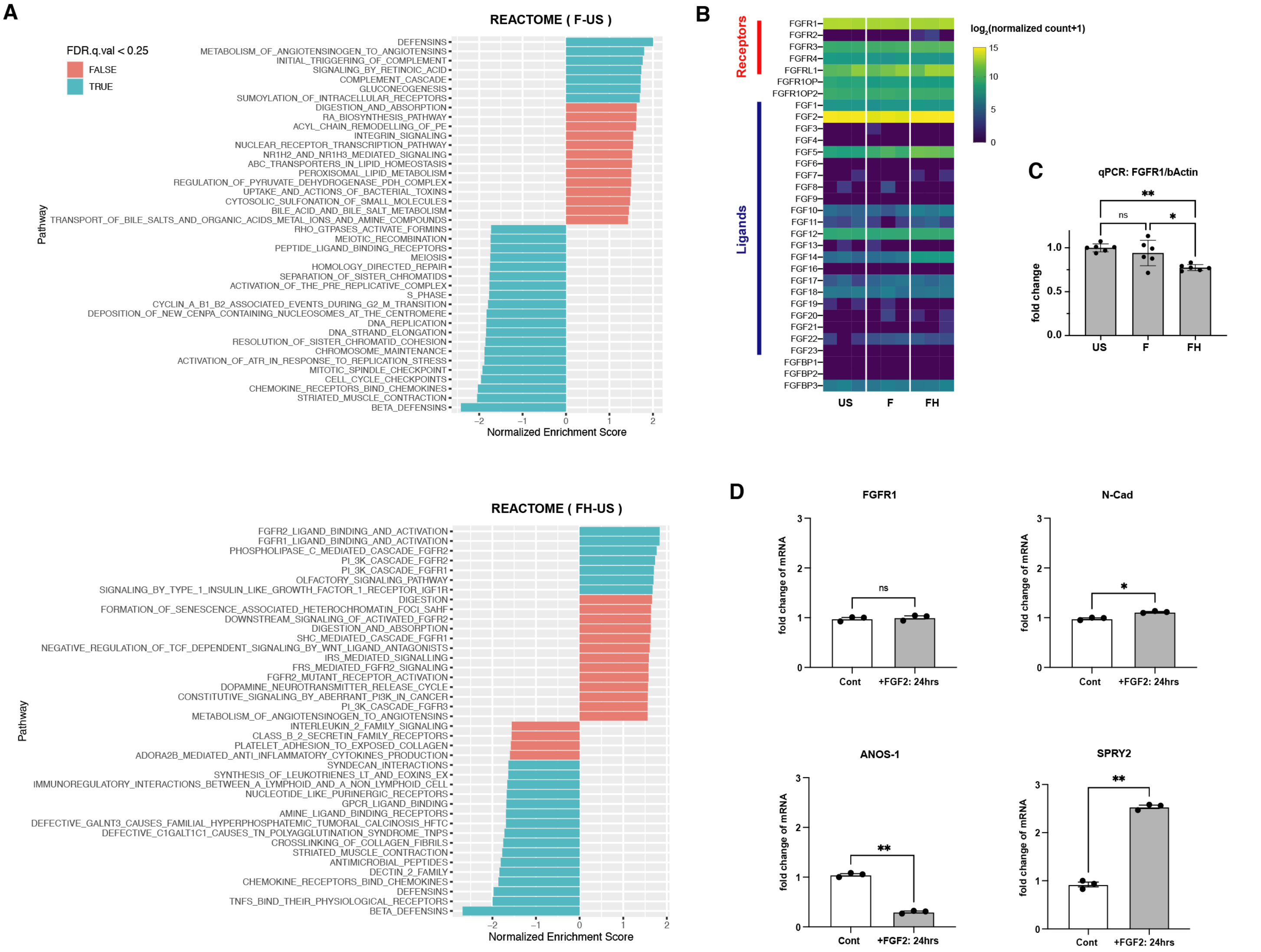


**Figure S2. Characterization of sorted cells and their effects on tumoroid size growth by VM formation.**

(A) Reactome Gene Set Enrichment Analysis (GSEA) result of F vs US comparison (top) and FH vs US comparison(bottom). Positive NES, enriched signature in F or FH; Negative NES, enriched signature in US.

(B) mRNA expression levels of FGFs, FGFRs and related genes selected from bulk RNA sequencing data. See the selected metadata Table S1.

(C) Relative expression of FGFR1genes quantified by real-time PCR (qPCR) in US, F, and FH cells (n=6, mean ± SEM). P-values were calculated using one-way ANOVA with Tukey's multiple comparison test.

(D) Quantification of the fold change in FGFR1, N-Cadherin, ANOS-1, and SPRY2 by FGF2 (100 ng/ml) treatment at the transcriptional level in US, acquired from RNA-seq data (n=3, mean ± SEM). P-values were calculated using two-tailed paired t-test.

#### Figure S3 related to Main Figure 4

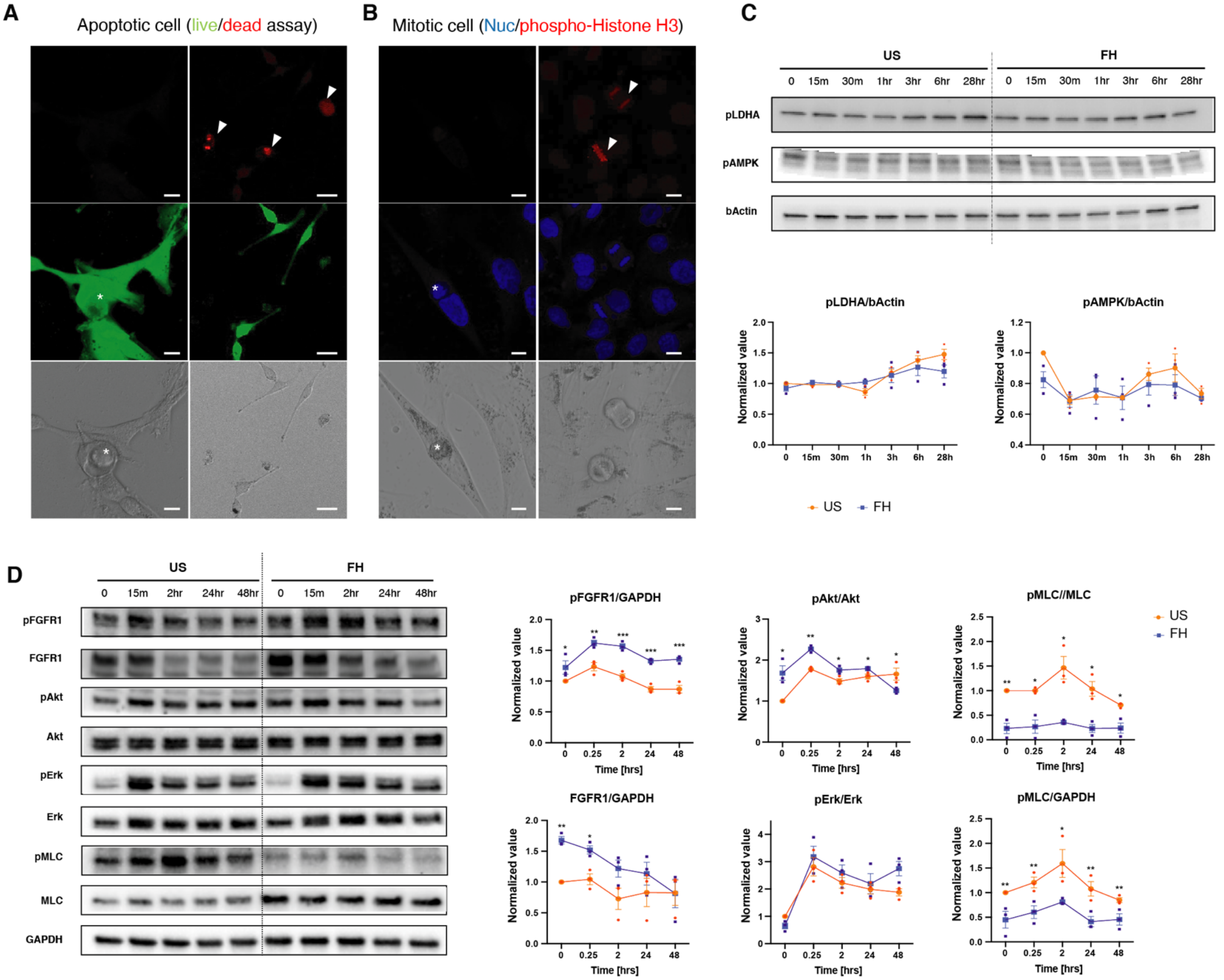


**Figure S3. Investigation of Entosis induction via FGF2-FGFR1 signaling.**

(A) Detection of the hosted cell's state using a live/dead assay. In dying and dead cells, a bright red fluorescence is generated upon binding to DNA. Asterisks (left) indicate a hosted cell, while arrowheads (right) indicate dying or dead cells not hosted by other cells. Scale bar (left): 20 $\mu$m; Scale bar (right): 100 $\mu$m.

(B) Detection of mitotic cells using phospho-histone H3 staining. Asterisks (left) indicate a hosted cell, while arrowheads (right) point to mitotic cells not hosted by other cells. Scale bar: 20 $\mu$m.

(C) LDHA and AMPK activity dynamics following FGF2 treatment in US and FH. β-Actin is used as a loading control (n=3, mean ± SEM). Adjusted p-values were calculated using unpaired multiple t-tests with two-stage linear step-up procedure of Benjamini, Krieger and Yekutieli.

(D) Representative immunoblots (left) showing temporal dynamics of downstream pathways of FGF2-FGFR1 in US and FH in response to FGF2 (100 ng/ml) treatment, with their quantification (right). GAPDH is used as a loading control (n=3, mean ± SEM). Adjusted p-values were calculated using unpaired multiple t-tests with two-stage linear step-up procedure of Benjamini, Krieger and Yekutieli.

#### Figure S4 related to Main Figure 5

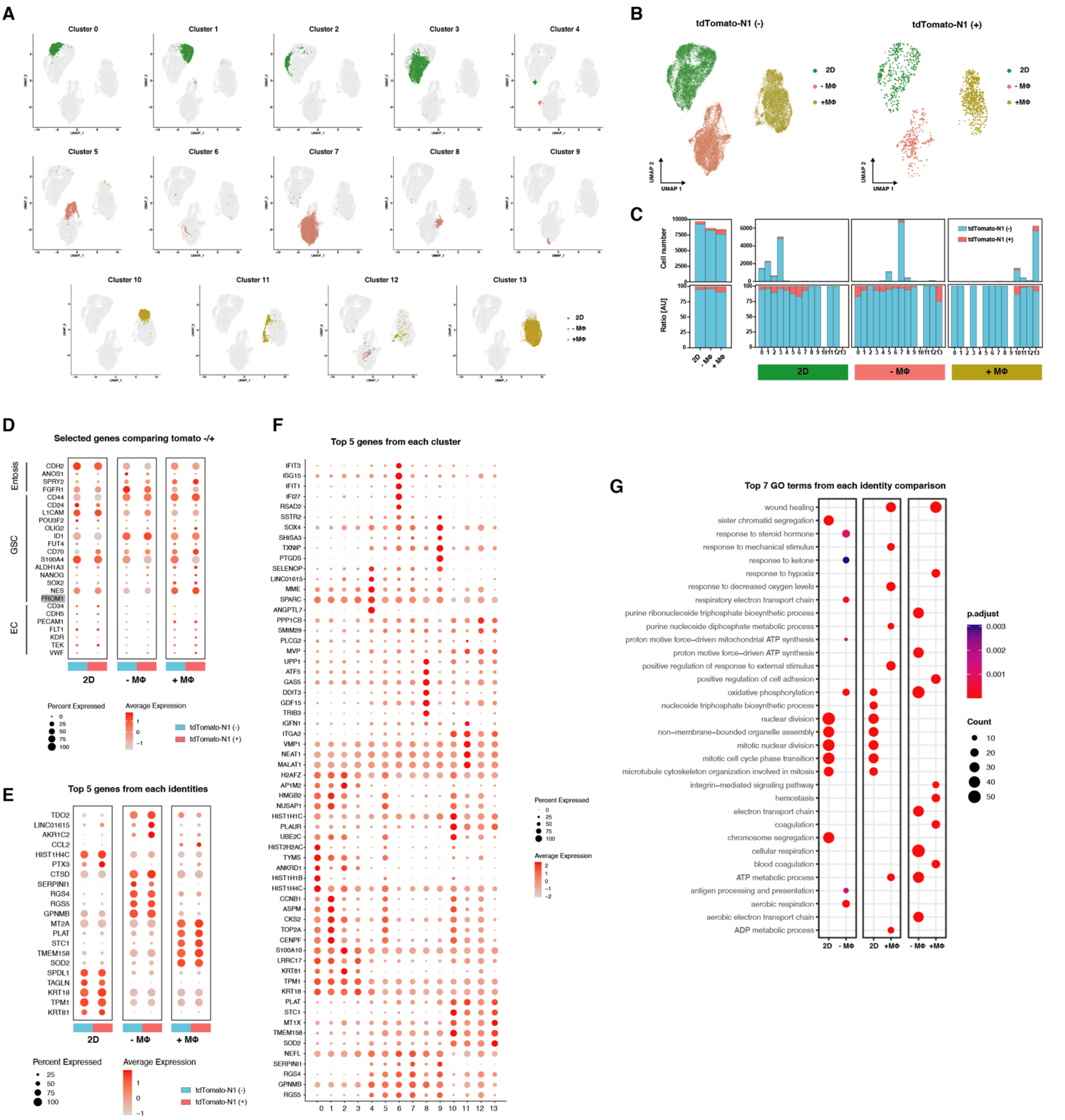


**Figure S4. GBM transcriptional changes and functional associations across culture conditions and clusters.**

(A) Individual clusters displayed on UMAP, color-coded by culture conditions.

(B) Separate UMAP visualizations of cells in which the tdTomato-N1 tagging gene was not detected (left) and was detected (right).

(C) Distribution cells in which the td-tomato tagging gene was not detected (blue) and was detected (red) across clusters.

(D) Dot plots displaying Entosis, glioblastoma stem cell (GSC) and endothelial cell-specific gene expression signatures in cells in which the td-tomato tagging gene was not detected (blue) and was detected (red) across culture conditions. Dot size represents the percentage of cells in which the gene of interest was detected and color indicates the gene's scaled average expression in those cells. PROM1 is shaded in gray to indicate that the gene was not detected.

(E) Dot plots displaying top 5 highly expressed genes in cells in which the td-tomato tagging gene was not detected (blue) and was detected (red) across culture conditions. Dot size represents the percentage of cells in which the gene of interest was detected and color indicates the gene's scaled average expression in those cells.

(F) Dot plots displaying top 5 highly expressed genes in each cluster. Dot size represents the percentage of cells in which the gene of interest was detected and color indicates the gene's scaled average expression in those cells.

(G) Top 7 Gene Ontology (GO) terms (BP: Biological Process) from comparisons between conditions. Cutoff: $p<0.05$; adjusted p (p.adj) $< 0.05$.

### Figure S5 related to Main Figure 5

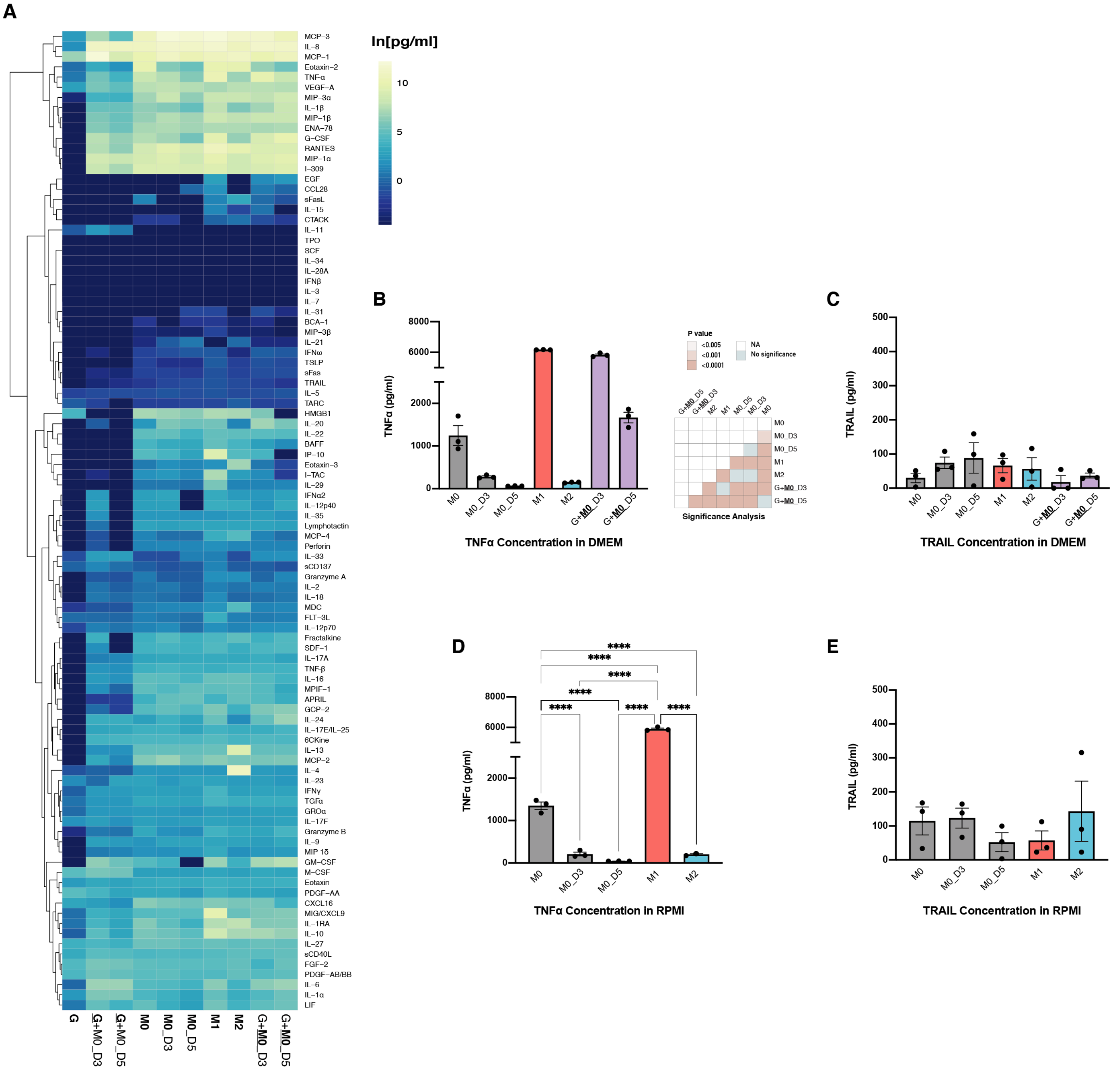


**Figure S5. Properties of macrophages differentially polarized under various conditions:** M0, newly differentiated macrophages ; M0_D3, M0 cultured for 3 days without additional treatment; M0_D5, M0 cultured for 5 days without additional treatment; M1, polarized by IFN-γ and LPS; M2, polarized by IL-4 and IL-13; G+**M0**_D3 , M0 co-cultured with GBM cells for 3 days; G+**M0**_D5 , M0 co-cultured with GBM cells for 5 days; A172 cells; macrophages, U937 cells differentiated using PMA.

(A) 96-plex secretion profiles from GBM cells and macrophages under various culture conditions: G, GBM cells only; **G**+M0_D3, GBM cells co-cultured for 3 days with M0 macrophages; **G**+M0_D5, GBM cells co-cultured for 5 days with M0 macrophages. DMEM was used as basal the medium for all conditions. See the full result in Table S2.

(B) ELISA analysis of TNFα and (C) TRAIL secreted from differentially polarized macrophages under various conditions (n=3, mean ± SEM): DMEM was used as basal the medium for all conditions. P-values were calculated using one-way ANOVA with Tukey's multiple comparison test.

(D) ELISA analysis of TNFα and (E) TRAIL secreted from differentially polarized macrophages under various conditions (n=3, mean ± SEM): RPMI was used as basal the medium for all conditions. P-values were calculated using one-way ANOVA with Tukey's multiple comparison test.

**Figure S6 related to Main Figure 7**

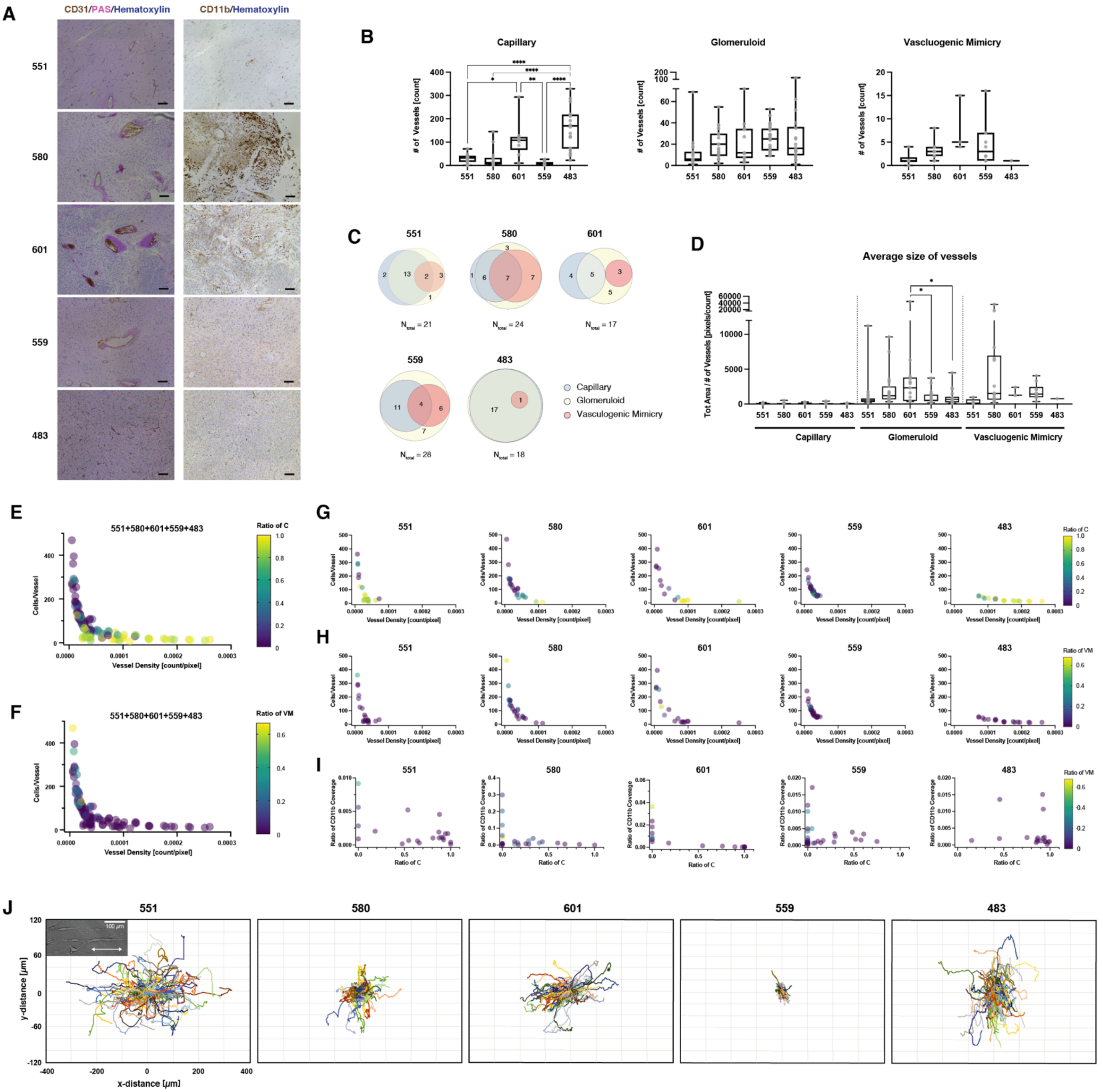

**Figure S6. Characteristics of vasculatures in GBM tissues from GBM patient tissues.**

(A) Representative histological images of GBM tissues from human patients (Fig. 1I) displaying vasculatures (CD31, PAS, and hematoxylin staining, top row) and distribution of monocytes or macrophages (CD11b and hematoxylin staining, bottom row). Scale bar: 100 $\mu$m

(B) Number of capillaries, glomeruloid, and vasculogenic mimicry vessels in tissues from human patients. Data are presented by box and whiskers with all individual points representing each image. P-values were calculated using one-way ANOVA with Tukey's multiple comparison test.

(C) Venn diagrams represent the count of images in which at least one of each vessel type was observed, out of the total number of images ($N_{tot}$) analyzed for each patient (bottom).

(D) Average size of vessels (measured in area) categorized by vessel types in tissues from human patients. Data are presented by box and whiskers with all individual points representing each image. P-values were calculated using one-way ANOVA with Tukey's multiple comparison test. pixel to distance ratio: 1.075 pixels/$\mu$m, pixel aspect ratio: 1.

(E) Scatterplot showing the correlation between vessel density and the number of cells supported by each vessel in each image corresponding to a data point. The color of each data point indicates the ratio of capillaries relative to all vasculature types in GBM tissues, including glomeruloid and vasculogenic mimicry vessels. Data from 551, 580, 601, 559, and 483 are integrated into the plot.

(F) Scatterplot showing the correlation as described in (E). The color of each data point indicates the ratio of vasculogenic mimicry vessels relative to all vasculature types in GBM tissues, including capillaries and glomeruloid vessels. Data from 551, 580, 601, 559, and 483 are integrated into the plot.

(G) Individual scatterplots for five patients showing the correlation as described in (E).

(H) Individual scatterplots for five patients showing the correlation as described in (F).

(I) Individual scatterplots for five patients showing the correlation between ratios of capillaries and ratio of CD11b coverages in each image corresponding to a data point. The color of each data point indicates the ratio of vasculogenic mimicry vessels relative to all vasculature types in GBM tissues.

(J) Dynamical tracking of the motility of individual patient derived cells on nanoscale ECM fiber-like ridges. A double-headed white arrow in the insect indicates the direction of ridges alignment.
