## Supplementary material for "Interplay between cytokine and FGF2 signaling in induction of entosis and vasculogenic mimicry response in glioblastoma": Methods

### EXPERIMENTAL MODEL AND STUDY PARTICIPANT DETAILS

**Collection approval by Yale and human patient specimen and information handling**
Institutional Review Board (IRB) approval and written informed consent was obtained at Yale New Haven Hospital for all patients included in this study. This study complied with all relevant ethical regulations. Patient data was stored and used anonymously in compliance with IRB and institutional regulations. All patients in this cohort underwent surgical resection. Surgical specimens were pathologically confirmed by a neuropathologist as World Health Organization (WHO) Grade 4 Glioblastoma.

**Extraction and isolation of GBM tissues and cells from patients**
All tumor samples were deidentified prior to use. The primary cultures were established as previously described[1,2]. Briefly, the tumor sample was washed in 1X RPMI, chopped into small pieces (<1 mm), and the fragments were digested with a mixture of collagenase type I and IV (0.5 mg/mL in PBS; Sigma-Aldrich, St. Louis, MO, USA) for 60 min at 37 °C. A single-cell suspension was prepared, washed and resuspended in DMEM/F12 supplemented with 20% fetal bovine serum (Life Technologies), in a humidified atmosphere of 5% CO2/air. Cells were split at no more than 1:3 per passage and used up to passage 7.

**Cell lines**
A172 (CRL-1620) cell line was purchased from ATCC. A172 cells were cultured in a growth medium of DMEM(Gibco) supplemented with 10% FBS (Life Technologies).
U-937(CRL-1593.2) cell line was purchased from ATCC. U-937 cells were cultured in a growth medium of RPMI-1640 (Gibco) supplemented with 10% FBS (Life Technologies).

**Primary endothelial cell**
GFP-expressing human brain endothelial cells (GFP-HBMEC) were purchased from Angio-Proteomie. GFP-HBMECs were cultured in a growth medium of M199 (Gibco) supplemented with 20% FBS (Life Technologies), 1% HEPES (Thermo Scientific), 1% Glutamax (Thermo Fisher), 1% antibiotic-antimycotic (Thermo Fisher), Heparin (25mg/500ml, Sigma Aldrich), and endothelial cell growth supplement (Sigma Aldrich, E2759) and used for further experiments at passage 9.

### METHOD DETAILS

**Formation of 3D tumoroids**
To produce uniformly-sized 3D tumorids from GBM cells cultured in 2D, we used microwells that are 800 μm in size (AggreWell-800 24 well plate, catalog # 34811, STEMCELL Technologie), following the manufacturer's protocol. To prevent cell attachment to the surface of the microwells, the plate was pre-treated with surfactant solution (Anti-Adherence Rinsing Solution, catalog # 07010, STEMCELL Technologie). Then, a cell suspension, with a concentration of $2 \times 10^5$ cells in 1.5ml medium, was added to each well, resulting in approximately 650-700 cells forming a tumoroid. The microplate was briefly centrifuged at $100 \times g$ to ensure cell capture in the microwells. Cells were incubated for 48 hours at 37°C with 5% $CO_2$ before use. The resulting tumoroid sizes ranged from approximately 200 μm to 300 μm.

**Development of 3D-GBM modeling micro-device (3D-GBM-MD): Fabrication, Cell Loading, and Culturing**
The 3D-GBM modeling micro-device (3D-GBM-MD) was designed and fabricated to mimic various aspects of the cellular composition and cell-cell interactions occurring in GBM. Detailed in Figure S1A,

the micro-device architecture consisted of two serial chambers separated by trapezoid pillars, enabling manipulation of the micro-environment elements in each chamber, while facilitating paracrine signaling across the entire structure. The device was prepared as detailed in our previous study[3]. The mold of this device was fabricated using stereolithography equipment (Protolabs Inc.) and a thermal-resistant resin. Components made of Polydimethylsiloxane (PDMS, Sylgard 184, Electron Microscopy Sciences) were then cast from this mold. Following a trimming process, the PDMS components were sterilized under UV light for a period exceeding one hour, after which they were placed on sterilized cover glasses. A fibrin gel solution was prepared by combining fibrinogen (Sigma Aldrich, F3879) at a concentration of 10 mg/ml with thrombin (Sigma Aldrich, T9549) at 5 U/ml in a 1:1 ratio. To this mixture, 0.15 U of aprotinin (Sigma Aldrich, SRE0050) was added per 1 ml of the solution[4]. All components were diluted in Dulbecco's Phosphate Buffered Saline (DPBS) without calcium and magnesium. GFP-HBMEC were suspended in this DPBS solution to achieve a final concentration of $2 \times 10^6$ cells/ml. Then the fibrin solution/HBMECs mixture was introduced into chamber C1. During this process, the pillars generated surface tension to prevent the gel-HBMEC mixture from seeping into the adjacent reservoir on the left side and chamber C2 on the right side. Next, a mixture of agarose (10mg/ml) and collagen type 1 (0.15mg/ml), along with GBM tumoroids—derived from patient cells or the GBM cell line A172—and macrophages differentiated from U937 cells, was labeled in green with 20 μM CellTracker™ Green CMFDA then introduced into chamber C2. 10-20 of tumoroid and $1 \times 10^5$ of macrophages/10 μl of gel was filled in chamber C2. GFP-HBMEC culture medium, either with or without 50 ng/ml of VEGF (Life Technologies, PHC9394), was added to the reservoir connected to chamber C1. Meanwhile, the culture medium for A172 cells was added to the reservoir connected to chamber C2.

**Immunofluorescence staining of cells cultured in 2D**

After fixing cells in 4% PFA for 30 min, cells were permeabilized with 0.1% Triton™ X-100 for 5 minutes, blocked with 10% goat serum at room temperature for 1 hour. Then cells were incubated with primary antibodies diluted in 10% goat serum at 4 °C overnight. After washing with DPBS three times, cells were incubated with secondary antibodies diluted in 10% goat serum for another 1 hour at room temperature. The nuclei were co-stained with Hoechst 33342. If necessary, F-actins were stained using approximately 66 nM of Alexa Fluor™ 647 Phalloidin, in combination with Hoechst 33342. After washing with DPBS three times, cell-in-cell formation was assessed using the transmission light detector then cells were imaged using fluorescent channels of a Leica SP8 confocal microscope.

Primary antibodies used: rabbit cathepsin B polyclonal antibody (dilution: 1 μg/ml, PA5-82618, Invitrogen); rabbit phospho-Histone H3 (Ser10) polyclonal antibody (dilution: 1/200, Cell Signaling, 9701); rabbit anti-beta catenin antibody (dilution: 2μg/ml, antibodies, A248301); mouse anti-N Cadherin (dilution: 5μg/ml, abcam, ab19348)

Secondary antibodies used: goat anti-rabbit IgG (H+L) Alexa Fluor™ 488 (dilution: 10 μg/ml Invitrogen, A-11008); goat anti-rabbit IgG (H+L) Alexa Fluor™ 594 (dilution: 10 μg/ml, Invitrogen, A-11012); goat anti-mouse IgG (H+L) Alexa Fluor™ 488 (dilution: 10 μg/ml, Invitrogen, A-11001); goat anti-mouse IgG (H+L) Alexa Fluor™ 594 (dilution: 10 μg/ml, Invitrogen, A-11032)

**Fluorescent imaging in 3D-GBM modeling micro-device (3D-GBM-MD)**

3D structures of vasculogenesis in the C1 chamber and VM formation within tumoroids in C2 chamber of the 3D-GBM-MD were captured using a confocal microscope (Leica, SP8, ×20 water immersion objective). This imaging was performed after fixing the cells in 4% PFA at 4 °C overnight, followed by permeabilization with 0.1% Triton™ X-100 for 1 hour. The F-actins and nuclei were then stained with approximately 66 nM of Alexa Fluor™ 647 Phalloidin and 1/200 of Hoechst 33342 diluted in PBS at 4 °C overnight and subsequently washed with PBS three times, each for 30 minutes. All solutions, including reagents and antibodies, were added to the reservoirs on both sides of 3D-GBM-MD. 3D images were reconstructed using IMARIS software (Bitplane), and the vessels in C1 were detected and quantified using the Surfaces function of IMARIS.

**Histology of tumoroids**

After fixing the cells in 4% PFA at 4 °C overnight, the bottom cover glass of the 3D-GBM-MD was removed. The tumoroids in C2 chambers were then extracted from the gel using sharp needles under a stereo microscope. Following three washes in DPBS, the tumoroids were incubated in 25% sucrose overnight at 4°C. They were subsequently embedded in Tissue-Tek O.C.T. Compound, stored at -80°C, and later processed for cryosectioning at a thickness of 14 µm. Cryosections were washed with DPBS then stained with periodic acid-Schiff (PAS) and counterstained with hematoxylin according to the manufacturer's protocol. After mounting with DPX mountant (Sigma Aldrich, 44581), sections were imaged using an optical microscope (Leica, Thunder Imager 3D Tissue).

For immunofluorescence staining, the cryosections were first blocked for 1 hour at room temperature in a blocking solution (1X DPBS, 10% goat serum, 0.1% Triton X-100). They were then incubated overnight at 4ºC with the primary antibody solution in 10% goat serum. This was followed by three washes in DPBS and a 1-hour incubation with the secondary antibody solution in 10% goat serum. After additional washes, the sections were mounted for imaging using Vectashield with DAPI (Vector laboratories, H-3500). All images were acquired using a Leica SP8 confocal microscope.

Primary antibodies used: rabbit anti-Laminin antibody (dilution: 1/50, abcam, ab11575); mouse anti-N Cadherin (dilution: 5µg/ml, abcam, ab19348); rabbit anti-VE-Cadherin (dilution: 1/200, Cell Signaling, 2500)

Secondary antibodies used: same as immunofluorescence staining of cells cultured in 2D.

**Histology of GBM tissues from human patients**

Formalin-fixed and Paraffin-embedded (FFPE) GBM tissue specimens were used for sectioning. FFPE blocks were trimmed, soaked in ice, and sectioned at 8µm using a microtome (Microm; Walidorf, Germany). Sections were floated on a water bath between 36-45 °C and then transferred onto adhesive-coated slides. Prepared slides were dried overnight at room temperature. Two adjacent sections per patient were acquired, with each section being used for different staining processes. One section underwent immunohistochemistry staining for rabbit anti-CD31 (abcam, ab28364) combined with periodic acid-Schiff (PAS)/hematoxylin, and the other for rabbit anti-CD11b (abcam, ab52478) with hematoxylin. The primary antibodies were detected using the rabbit-specific HRP/DAB (ABC) Detection IHC Kit (abcam, ab64261), following the manufacturer's protocol. After mounting with DPX mountant (Sigma Aldrich, 44581), the entire area of each section was scanned using an optical microscope (Leica, Thunder Imager 3D Tissue, 10x objective) then automatically stitched as a one image per section.

**Quantification of histology of GBM tissues from human patients**

Each section, stained for CD31/periodic acid-Schiff (PAS)/hematoxylin and for CD11b/hematoxylin, from the same patient was loaded into ImageJ, then stacked and aligned using the Registration plugin with Linear stack alignment with SIFT (Transformation: Similarity, maximal alignment error: 100, which needed to be adjusted for the best fit). Multiple images, including a local area (1300 pixels by 1000 pixels, equivalent to the size imaged by a 10x objective lens), were then selected and extracted to cover the whole area without overlapping. Each local image stack was then divided again into CD31/periodic acid-Schiff (PAS)/hematoxylin staining and CD11b/hematoxylin staining for further analysis to quantify vessel density, cell density, PAS-positive area, and CD11b-positive area. The code for the quantification in ImageJ was provided. Manual selection of vessels was required in the quantification. Vessels were categorized into capillary, glomeruloid vessels, and vasculogenic mimicry (VM) vessels based on their morphologies as depicted in Fig. 1K: Capillaries had a very thin wall, typically just one cell thick, with a continuous lining of CD31-positive endothelial cells and with no significant PAS-positive ECM deposition around. Glomeruloid vessels displayed a complex, glomerulus-like structure, dilated, discontinuous lining of CD31-positive endothelial cells, and thick PAS-positive ECM deposition around. VM vessels displayed a dilated and bare lumen without CD31-positive endothelial cells, and thick PAS-positive ECM deposition around. Vessels that underwent thrombosis or were indistinguishable from arterioles/venules, which were rarely observed in the sections, were counted in the total number of vessels but were not categorized into a specific vessel type. The threshold values for PAS and CD11b positive area were set to distinguish clear deposition of PAS-positive ECM around glomeruloid vessels and VM vessels, and rounded CD11b-positive monocytes from unspecific binding to the background.

**Tracking tumoroids size growth**

The growth of tumoroids in C2 was monitored using a phase contrast microscope (ZEISS, Axiovert 25, ×10 objective) on day 0, day 3, and day 7. The entire area of C2 was captured in multiple images, which were then stitched together for analysis. These images were converted into binary format, where black objects represent tumoroids, while small dots indicative of macrophages were excluded. The sizes of the tumoroids were measured using ImageJ, and each tumoroid was identified by its location, with sizes normalized to their initial measurements on day 0. Three 3D-GBM-MD were used for each condition and acquired data from the three chips were combined in the plot.

**FITC-Dextran transport test**

After imaging the tumoroids stained for F-actins and nuclei, a 70 kDa FITC-dextran solution at a concentration of 30 µg/ml in PBS was added to both reservoirs of the 3D-GBM-MD. The chip was then incubated for 30 minutes prior to further imaging at room temperature. After 1 hour and again at 24 hours, the tumoroids exhibiting VM were imaged using a confocal microscope (Leica, SP8, ×20 water immersion objective). The intensity of the FITC signal was analyzed using ImageJ.

**Hypoxic core detection**

To detect the hypoxic core in tumoroids cultured in the 3D-GBM-MD, we used the Hypoxyprobe™ Plus kit (HPI, HP2-100), following the manufacturer's protocol. The culture medium, containing 400 µM of Hypoxyprobe™-1 (pimonidazole hydrochloride), was added to the chip's reservoirs and incubated for 2 hours. Then, the tumoroids were fixed, extracted, sectioned, and prepared for staining as described in the 'Histology of tumoroids.' Sections were incubated with FITC-conjugated mouse anti-pimonidazole monoclonal antibody (FITC-Mab1, 1/100 dilution). After washing, the sections were mounted for imaging using Vectashield with DAPI (Vector laboratories, H-3500). All images were acquired using a Leica SP8 confocal microscope.

**MRI Image Analysis**

Pre-operative Magnetic Resonance Imaging (MRI) was performed on patients with GBM included in this study. T1-weighted (T1), T1-weighted image with contrast (T1+C), T2-weighted (T2), fluid attenuated inversion recovery (FLAIR), diffusion weighted imaging (DWI), and apparent diffusion coefficient (ADC) were analyzed. The following features were characterized: tumor location (e.g. frontal, parietal, occipital, temporal), contrast-enhancement (e.g. diffuse, ring-enhancing, scattered), T2/FLAIR signal, and intratumoral diffusion restriction (hyperintense on DWI, hypointense on ADC). Minimum, maximum, and mean ADC values were calculated by performing a 3-dimensional region of interest (ROI) encompassing the entirety of the tumor on x, y, and z planes on ADC images.

**Cell sorting via chemotaxis**

We sorted the cells based on their chemotactic capacity in FGF2 gradients under different conditions. The microfluidic chip used for these experiments was designed with two reservoirs (dimensions: 30 mm by 15 mm, 7.2 mm height) and 15 channels, each 2 mm long, connecting them. The channels have a width of 500 µm and a height of 200 µm. The mold for this chip was created using a 3D printer (Ultimaker 2) with PLA filaments. Components made of Polydimethylsiloxane (PDMS, Sylgard 184, Electron Microscopy Sciences) were cast from this mold. After a trimming process, the bottom surface of the PDMS components and cover glasses were treated with a corona discharge (Electro-Technic Products, BD-20AC) and then bonded together. The assembled chips were sterilized under UV light for over an hour. After filling the reservoirs and channels with 70% ethanol, washing them three times with DPBS, and completely aspirating the liquids, each channel was filled with a 0.4 mg/ml collagen type 1 solution. Excess collagen solution was carefully aspirated, and the chip was then incubated at 37°C for 30 minutes. The surfaces of the reservoirs were coated with 2 μg/ml of laminin for 2 hours, after which A72 cells were seeded to cover 70% of the surface of the left reservoir. Both reservoirs were filled with an equal amount of culture medium, and the chip was placed at 37°C for one day to allow cell adherence. Subsequently, FGF2 was added to the right reservoir to achieve a final concentration of 200 ng/ml, with an equivalent volume of medium added to the left reservoir to even out the levels. The medium and FGF2 were refreshed daily for 3 days under normoxic conditions at 37°C, which facilitated cell migration toward the right reservoir via chemotaxis in response to the FGF2 gradient. Then, cells from the left and right reservoirs were collected separately by trypsinization. For hypoxic conditions, chips were placed in a modular incubator chamber (Billups-Rothenberg Inc, MIC-101) filled with 1% oxygen at 37°C. Similar to the normoxic condition, the medium and FGF2 were refreshed, and 1% oxygen gas was refilled daily. After 7 days, cells from the left and right reservoirs were collected. Fewer than 50 cells were collected from each chip. These collected cells were propagated through subculture for a month and then cryopreserved in multiple vials for future experiments.

**Cell-in-cell assessment**

Two sets of unsorted A172 cells were pre-labeled for dual-color imaging: one set with 20 μM CellTracker™ Green CMFDA and the other with 20 μM CellTracker™ Orange CMTMR, each incubated for 45 minutes at 37°C. Sorted, either under normoxic or hypoxic conditions, A172 cells were also labeled with 20 μM CellTracker Green CMFDA under identical conditions, enabling clear differentiation between red-labeled and green-labeled cells in co-culture experiments. After co-plating either a total of $1 \times 10^4$ red(unsorted)-green(unsorted)cells or red(unsorted)-green(sorted) cells at a 1:1 ratio in each well of an 8-well chambered coverglass (Nunc™ Lab-Tek™ II, 155409), precoated with 2 μg/ml of laminin for 24 hours, the cells were incubated for an initial 24 hours. This was followed by another 24-hour period with or without treatment of 100 ng/ml FGF2 (10781-722, VWR), 0.5 μM of Y-27632 (688001, Millipore Sigma), or 20 μM of ADH-1 (29981, Cayman). As a control for the Y-27632 and ADH-1 treatments, the culture medium was supplemented with 0.5% DMSO, the same amount used for dissolving these drugs. Cells were imaged using a confocal microscope (Leica, SP8, with a ×20 water immersion objective) using filter sets for green (Ex=492 nm, Em=517 nm) and red (Ex=541 nm, Em=565 nm) fluorescence. Z-stacks were acquired to assess cell-in-cell formation, where one cell is entirely engulfed by another. Three images from different positions were analyzed to count cell-in-cell formation in each group, with each image containing approximately 100 to 300 cells. Cell-in-cell formations were distinguished by the differing colors of the host and hosted cells; structures with the same colors were not considered in data. The cell-in-cell ratio was calculated based on the number of red cells hosting green cells relative to the total number of red cells, and similarly for green cells hosting red cells relative to the total number of green cells.

**Bulk RNA sequencing analysis of unsorted and sorted A172 cells**

Total cellular RNA from cells using the RNeasy Plus Mini Kit (Qigen, 74134) as per the manufacturer's instructions. Three biological replicates per sample were sent to the Yale Center for Genome Analysis (YCGA) for cDNA library preparation and sequencing to HiSeq paired-end 100 bp on Illumina NovaSeq, targeting 25 million read pairs, using YCGA's standard alignment pipeline. Reads were aligned to the hg38 genome using the STAR aligner (v 5.0.0). YCGA provided normalized counts and differentially expressed genes (DEGs; adjusted p-value < 0.05 and |log2FC| > 1) computed using the DESeq2 package in R. edgeR and ggplot2 were used to visualize normalized counts by clusters with a heatmap, and to identify variance among samples using principal component analysis (PCA). Gene-set enrichment analysis (GSEA) was performed with GSEA 4.1.0 from UCSD/Broad Institute using the REACTOME database. Gene Ontology (GO) biological process enrichment analysis was performed with lists of up-regulated or down-regulated genes (p-value < 0.05 and |log2FC| > 1) on Enrichr, a web-based and mobile software application.

**Immunoblotting**

Cells prepared for immunoblotting were washed with DPBS washing ($MgCl_2$ 1 mM, $CaCl_2$ 1 mM) then lysed with RIPA buffer mixed with Halt proteases and phosphatase inhibitor cocktail (Thermo Scientific), following the general protocol for western blotting of Bio-Rad. After measuring protein concentrations with BCA assay kit, they were equalized then followed by a denaturation in 4x Laemmli buffer mixed with 2-mercaptoethanol by boiling at 95 °C for 5 min. Then the lysates were loaded to 4-20% Mini-PROTEAN TGX precast gels (Biorad) for electrophoresis and proteins were transferred to 0.2 mm nitrocellulose membrane using Tran-Blot Turbo (Biorad) set at 2.5V, 25 A for 10 minutes. The membrane was blocked with 3% BSA in TBST and incubated with primary antibodies overnight at 4 °C and followed by horseradish-peroxidase-coupled secondary antibody for 1 hour at room temperature. Between each step, the membrane was washed three times with TBST for 15 min each. Finally, the membrane was incubated with Clarity Western ECL (Bio-Rad) substrates and bands were imaged using ChemiDoc XRS (Bio-Rad). The original images without changing the grayscale curve were used for intensity measurement in ImageJ. All immunoblotting images presented in the figures are the original images. Each band was normalized to GAPDH or ß-actin expressions and then further normalized to the experimental control.
To evaluate the activation of downstream pathways of FGFR1-FGF2 (Figure. 4A, S3D), 100 ng/ml of FGF2 was added to indicated cells, and then samples were harvested at following time points after the treatment: 15 min, 2 hours, 24 hours, 48 hours.
To inhibit phosphorylation of FGFR1, the most upstream component of the FGFR1-FGF2 pathway (Figure 4B), Unsorted A172 cells were treated with 0.1 μM of PD 173074 (Selleckchem, S1264) for 24 hours. To perturb the signaling pathways, cells were treated with 1 μM of SB 590885 (Cayman, 16643) and 2 μM of wortmannin (Cayman, 10010591), either separately or in combination, as indicated. For the control, the culture medium was supplemented with 0.1% DMSO, the solvent used for these drugs.
All siRNA experiments were performed at a final concentration of 10 nM siRNAs for control (Silencer, Cat # AM4611, Thermo Fisher) , FGFR1 (Silencer, Assay ID:1216, Thermo Fisher), N-Cadherin (Silencer, Assay ID: 145997, Thermo Fisher), 20 nM for SPRY2 (Silencer, Assay ID: 107851, Thermo Fisher), and 30 nM for ANOS-1 (Silencer, Assay ID: 8149, Thermo Fisher) using Lipofectamine RNAiMAX Transfection Reagent (13778075, Invitrogen), according to the manufacturer's instructions. At 48 hours after siRNA transfection, protein knockdown was assessed by western blot analysis.

Primary antibodies used: rabbit cathepsin B polyclonal antibody (dilution: 1 μg/ml, PA5-82618, Invitrogen);
pFGFR1 (dilution: 0.2 μg/mL, Cat # 06-1433, Millipore Sigma); FGFR1 (dilution: 1/1000, Cat # 9740, Cell Signaling); pAkt (dilution: 1/1000, Cat #4060, Cell Signaling); Akt(dilution: 1/1000, Cat #9272, Cell Signaling); pErk (dilution: 1/1000, Cat #9101, Cell Signaling); Erk(dilution: 1/1000, Cat #9102, Cell Signaling); pMLC2(dilution: 1/1000, Cat #3674, Cell Signaling); MLC2(dilution: 1/1000, Cat #8505, Cell Signaling); SPRY2(dilution: 1/1000, Cat #14954, Cell Signaling), GAPDH(dilution: 1/1000, Cat #2118, Cell Signaling), β-Actin (dilution: 1/1000, Cat #3700, Cell Signaling)
Secondary antibodies used: ECL Rabbit IgG, HRP-linked whole Ab from donkey (dilution: 1/5000, NA934, GE HealthCare); ECL Mouse IgG, HRP-linked whole Ab from sheep (dilution: 1/5000, NA931, GE HealthCare)

**Co-immunoprecipitation**
Co-immunoprecipitation experiments with A172 cell lysates were performed using the Pierce Co-Immunoprecipitation (Co-IP) Kit (Thermo Scientific, 26149), according to the manufacturer's recommendations. To briefly note the important values in the protocol, 50 μl of AminoLink Plus Coupling Resin was added into a Pierce Spin Column and washed. Separately, 3 μg of IgG (H2113) and 3 μg of FGFR1 antibodies were prepared, each adjusting the volume to 200 μl with 1X Coupling Buffer. Each antibody was then added to its respective column and incubated on a rotator at room temperature for 2 hours before washing. 2 mg of cell lysates, pre-cleared using control agarose resin, was added to each antibody-coupled resin and incubated at 4 °C for 1 hour with gentle end-over-end mixing, followed by centrifugation.

After eluting proteins with 50 µl of Elution Buffer at room temperature, the samples were then denatured and processed according to the immunoblotting protocol.

**Quantitative Real-Time PCR**

RNAs were purified from indicated cells using the RNeasy Plus Mini kit (QIAGEN), adhering to the guidelines provided by the manufacturer. For the reverse transcription process, iScript Reverse Transcription Supermix (Bio-Rad) was employed, following the steps: initial priming at 25 °C for 5 minutes, reverse transcription at 46 °C for 20 minutes, and inactivation at 95 °C for 1 minute, using a cycler (C1000 Touch, Bio-Rad). Gene expression analysis was conducted using TaqMan Fast Advanced Master Mix (Applied Biosystems), following the protocol of 2 minutes incubation at 50 °C, 20 seconds of polymerase activation at 95 °C, and 40 PCR cycles consisting of 5 seconds of denaturing at 95 °C and 30 seconds of annealing/extending at 55 °C, using a real-time PCR system (CFX384 Touch, Bio-Rad). The FGFR1(primer Assay ID: Hs00241111_m1, FAM-MGB, Thermo Fisher) results were normalized to ß-actin levels (primer Assay ID: Hs01060665_g1, VIC-MGB_PL, Thermo Fisher) from the corresponding wells. All experiments were replicated three times, with each set of replicates derived from three separate biological samples.

**Live/Dead Cell assay**

To discriminate between live and dying/dead cells based on intracellular esterase activity and plasma membrane integrity, we used the LIVE/DEAD™ Cell Imaging Kit (Invitrogen, R37601), following the manufacturer's protocol. After incubating the cells with reagents for 15 minutes, we imaged them using filter sets for green (live cells, Ex=488 nm, Em=515 nm) and red (dying/dead cells, Ex=570 nm, Em=602 nm) fluorescence with a confocal microscope (Leica, SP8, ×40 oil immersion objective).

**Cell culture for death detection in cell-in-cell by TUNEL reaction and Cathepsin B staining**

After co-plating a total of $1 \times 10^4$ unsorted and sorted, under hypoxic condition, A172 cells at a 1:1 ratio in each well of an 8-well chambered coverglass (Nunc™ Lab-Tek™ II, 155409), precoated with 2 µg/ml of laminin for 24 hours, the cells were incubated for 24 hours in fresh medium, following another 24 hours in either fresh medium or conditioned medium from a 2-day co-culture of M0-state U937 macrophages and A172 cells.

**Death detection in cell-in-cell by TUNEL reaction**

To detect apoptotic host or hosted cells, we used the In Situ Cell Death Detection Kit, TMR red (Roche, 12156792910), following the manufacturer's protocol for adherent cells, and co-stained the cells with Hoechst 33342. Cell-in-cell formation was assessed using the transmission light detector of a confocal microscope (Leica, SP8, ×40 oil immersion objective). TMR red and Hoechst 33342 signals in the nuclei of host and hosted cells were detected using fluorescent channels. TUNEL-positive host or hosted cells were counted regardless of intensity, while the excitation light intensity and emission light gain were set at levels to minimize false signals and background noise, ensuing accurate identification without overexciting non-apoptotic cells.

**Acridine Orange (AO) staining and quantification**

After co-plating unsorted and sorted, under hypoxic condition, A172 cells at a 1:1 ratio in each well of an 8-well chambered coverglass (Nunc™ Lab-Tek™ II, 155409), precoated with 2 µg/ml of laminin for 24 hours, the cells were incubated for 24 hours, following another 24 hours in various media: fresh medium; medium containing 10 µg/ml or 100 µg/ml of TNF-α; medium with 10 µg/ml or 100 µg/ml of TRAIL; or conditioned medium from a 2-day co-culture of M0-state U937 macrophages and A172 cells. After aspirating the medium, 450 µl of blank medium were added to each well. The Acridine Orange staining solution (Immunochemistry Technologies, 6130) was then diluted at a 1:20 ratio in PBS, and 50 µl of this diluted solution was added to each well, equal to a final concentration of 5.0 µM. Following a 25-minute incubation at 37°C, the cells were washed with PBS and replenished with fresh blank medium.

Subsequently, the cells were immediately analyzed using a confocal microscope (Leica, SP8, ×40 oil immersion objective). Cell-in-cell formation was assessed using the transmission light detector and then imaged with filter sets for green (Ex=492 nm, Em=540 nm) and red (Ex=540 nm, Em=650 nm) fluorescence. The fluorescent signal was quantified as the ratio of the 'red' over 'green' signal in the area of the entotic vacuoles surrounding the hosted cells or in the cytosol of host cells, excluding the nucleus and hosted cell.

**Collecting supernatant for multiplex analysis and ELISA**

$1 \times 10^6$ U937 cells were seeded into each well of a 12-well plate containing RPMI-based culture medium. They were then differentiated into M0 macrophages by treating them with 200 nM PMA (Phorbol 12-myristate 13-acetate) for 3 days. After aspirating the medium, 1.5 ml of DMEM supplemented with 10% FBS or RPMI supplemented with 10% FBS was added to each well. Following 24 hours of incubation, the medium from each well was collected into a 15 ml tube and centrifuged at 900 rpm. Subsequently, 1 ml of the supernatant was collected for analyzing secretions from M0 macrophages. The supernatant was collected at three time points: immediately after differentiation to day 1 (M0), from day 2 to day 3 (M0_D3), and from day 4 to day 5 (M0_D5). M1 macrophages were induced from M0 macrophages by treatment with 200 ng/ml IFN-γ and 100 ng/ml LPS, while M2 macrophages were induced using 20 ng/ml of IL-4 and 20 ng/ml of IL-13. Both treatments were conducted for 24 hours in 1.5 ml of either DMEM or RPMI supplemented with 10% FBS. Supernatants from M1 and M2 were collected directly as described above, without changing the medium. It is important to note that each supernatant includes IFN-γ and LPS, or IL-4 and IL-13. For co-culture with tumoroids made of A172 cells, 133 tumoroids were seeded with an agarose(10mg/ml)/collagen type 1(0.15mg/ml) mixture into the upper side of a 0.4 μm pore polyester the membrane insert (Corning, 3460), which was then transferred to each well containing M0 macrophages at the bottom. For the co-culture, cells were incubated in DMEM-based medium. For the GBM-only group, the transwell insert was transferred to a blank well and incubated with 0.5 ml of fresh medium for 24 hours. Transwell inserts from co-cultured samples were moved to a new blank well plate on days 2 (**G**+M0_D3) and 4 (**G**+M0_D5) and incubated with fresh medium for 24 hours. Additionally, 1.5 ml of fresh medium was added to the remaining macrophages in the 12-well plate on days 2 (G+**M0**_D3) and 4(G+**M0**_D3), followed by a 24-hour incubation. Supernatants were then collected as described above. Triplicate samples were prepared for each group. For Multiplex analysis, triplicate samples were mixed at the same ratio to average the biological replicates, whereas for ELISA, each sample was analyzed separately.

**Multiplex analysis of cytokines**

This study used Luminex xMAP technology for multiplexed quantification of 96 Human cytokines, chemokines, and growth factors. The multiplexing analysis was performed using the Luminex™ 200 system (Luminex, Austin, TX, USA) by Eve Technologies Corp. (Calgary, Alberta). Ninety-six markers were simultaneously measured in the samples using Eve Technologies' Human Cytokine 96-Plex Discovery Assay® which consists of two separate kits; the Panel A 48-plex and the Panel B 48-plex (MilliporeSigma, Burlington, Massachusetts, USA). The assay was ran according to the manufacturer's protocol. The Panel A 48-plex consisted of sCD40L, EGF, Eotaxin, FGF-2, FLT-3 Ligand, Fractalkine, G-CSF, GM-CSF, GROα, IFN-α2, IFN-γ, IL-1α, IL-1β, IL-1RA, IL-2, IL-3, IL-4, IL-5, IL-6, IL-7, IL-8, IL-9, IL-10, IL-12(p40), IL-12(p70), IL-13, IL-15, IL-17A, IL-17E/IL-25, IL-17F, IL-18, IL-22, IL-27, IP-10, MCP-1, MCP-3, M-CSF, MDC, MIG/CXCL9, MIP-1α, MIP-1β, PDGF-AA, PDGF-AB/BB, RANTES, TGFα, TNF-α, TNF-β, and VEGF-A. The Panel B 48-plex consisted of 6CKine, APRIL, BAFF, BCA-1, CCL28, CTACK, CXCL16, ENA-78, Eotaxin-2, Eotaxin-3, GCP-2, Granzyme A, Granzyme B, HMGB1, I-309, I-TAC, IFNβ, IFNω, IL-11, IL-16, IL-20, IL-21, IL-23, IL-24, IL-28A, IL-29, IL-31, IL-33, IL-34, IL-35, LIF, Lymphotactin, MCP-2, MCP-4, MIP-1δ, MIP-3α, MIP-3β, MPIF-1, Perforin, sCD137, SCF, SDF-1, sFAS, sFASL, TARC, TPO, TRAIL, and TSLP.

Blank culture medium was analyzed alongside the supernatants from A172 cells and U937 macrophages, which are specified above. Although the values obtained from the blank medium were near zero, all data were subtracted by the values from the blank medium to account for any baseline factors in the culture medium. For data analysis purposes, any final values out of range (OCR) were systematically replaced with the lowest or highest value within the range. The data were then natural log-transformed. Samples were visualized and hierarchically clustered using the heatmap.2 and function from gplots and pheatmap packages library in R.

**ELISA for TNFα/TRAIL**

To measure secretion from glioblastoma cells and macrophages under different culture conditions, the supernatants (three biological replicates per condition) mentioned above were diluted at a 1:5 ratio. Subsequently, the secreted levels were measured using enzyme-linked immunosorbent assay (ELISA) kits, following the manufacturer's recommendations, for TNFα (Invitrogen, Cat No. 88-7346-88) and TRAIL (R&D Systems, Cat No. DTRL00).

**Generation of fluorescent-expressing A172**

tdTomato-N1 was a gift from Michael Davidson, Nathan Shaner, and Roger Tsien (Addgene plasmid # 54642). The QIAprep Spin Miniprep kit (QIAGEN, Cat No. Q27106,) was used to purify plasmids. tdTomato was transfected into A172 cells at 70% confluency in a 10cm dish using the Lipofectamine™ 3000 system. We diluted 41μl of Lipofectamine™ 3000 reagent (Thermo Fisher, Cat No. L300000 in 1.5ml Opti-MEM. The master mix of tdTomato plasmid at 5μg/μl was also diluted in 1.5ml Opti-MEM with 35μl of P3000™ Reagent. We then added the master mix to the Lipofectamine™ 3000 reagent in a 1:1 ratio and incubated it for 15 minutes at room temperature. The cells were incubated for 48 hours at 37°C. Subsequently, the fluorescence was confirmed under a microscope. G418 (IBI Scientific, Cat No. 75794-854) was used to select the cells, tested at concentrations of 100μg/ml, 200μg/ml, 300μg/ml, 400μg/ml, and 500μg/ml. To establish a stable fluorescent cell line, a cell suspension solution (5 cells/ml), which had been passed through a 40μm cell strainer mesh (Fisher Scientific, Cat No. 352340) to avoid any cell clumps, was transferred to a 96-well plate labeled with singlets. After a week, the plates were scanned to identify the growth of colonies expressing tdTomato under a fluorescent microscope (Zeiss, Axiovert 25). During this process, the G418 concentration was reduced to 200μg/ml to create a less harsh environment for the colonies. For further expansion, 400μg/ml of G418 was used, but G418 was not added in the process of preparing cells for scRNA sequencing.

**scRNA sequencing analysis of unsorted and sorted cells mixture cultured under different conditions**

Unsorted and sorted, under hypoxic condition, A172 cells were mixed at a 1:1 ratio and then cultured in 2D, or in a 3D agarose/collagen mixture gel, in the form of tumoroids, either without (-MΦ) or with macrophages (+MΦ). The ratio of tumoroids to macrophages was maintained consistently for supernatant collection for multiplex analysis and ELISA. The tumoroids encapsulated in the gel were placed in the upper insert, and the macrophages were placed in the bottom well of a 75 mm transwell with a 0.4 μm pore polycarbonate membrane insert (Corning, 7910), as depicted in Figure 5A. The number of cells per tumoroid and the ratio of tumoroids to macrophages in a well were set to be the same as in the 3D-GBM-MD. A DMEM-based culture medium for A172 cells was used for 3 days. Then, the tumoroids in the upper insert were extracted from the gel using sharp needles under a stereo microscope. Following a wash in DPBS, the tumoroids were dissociated with a 0.25% trypsin, 0.03% EDTA solution (Life Technologies), and then the cells were filtered through 70 μm cell strainers twice to remove debris from the gel.

Single-cell suspensions in DPBS were sent to the Yale Center for Genome Analysis (YCGA) for library preparation with 10x Genomics Chromium Single Cell 3' Reagent Kits v.3.1, and 10,000 cells per sample were sequenced separately on a NovaSeq6000 at the YCGA, using YCGA's standard alignment pipeline. Reads were aligned to the hg38 genome using the STAR aligner (v 5.0.0) with a modified reference including the tdTomato-N1 sequence (available at www.addgene.org/54642). Downstream analyses of gene expression matrices were processed in R using the Seurat package. Input data from the three separate sequences for each condition were loaded together as a row count, each labeled with its respective condition. Higher-quality cells within the range of the following criteria remained, and rRNA genes were excluded: 200 < nFeature < 6000, 200 < nCount < 30000, and percentage of mitochondrial genes < 25%; genes were detected in at least 10 cells. 9,683, 8,586, and 8,394 of the cells were kept in the analyses for 2D, -MΦ, and +MΦ, respectively.

Data were then normalized with SCTransform. Cells were labeled as '_tomato' after the original labeling if they expressed the tdTomato-N1 gene, regardless of the expression level. PCA (npc=100, with all 21839 features) and nonlinear dimensional reduction (UMAP) were obtained using RunPCA and RunUMAP function, respectively. FindNeighbors function was performed using PCA dimensions [1:100] with Shared Nearest Neighbor (SNN) setting and FindClusters was performed with resolution 0.4.

FindAllMarkers function was used to find genes that are distinctly expressed in each cluster or in each identity by switching the active identity by Idents function.

The top 300 differentially expressed genes in the comparison of each identity or cluster were used to perform GO biological process enrichment analysis using the enrichGO function. GSEA was performed with gseaKEGG function run on KEGG database. The pairwise similarity between GSEA terms was calculated using the pairwise_termsim function. The results were visualized using the emapplot function, which displays the similarity among terms. Additionally, the connections between terms and genes were illustrated using the cnetplot function. Gene sets related to entosis, glioblastoma stem cell (GSC), endothelial cell (EC), and predefined gene sets of specific GSEA or GO terms were displayed using the DotPlot function. Scores for each cell were calculated using the AddModuleScore function based on the expression of the genes in each term and mapped onto UMAP plots. The terms of interest and their corresponding IDs are as follows: Cell Migration - GO:0016477; ECM-Receptor Interaction - hsa04512; Extracellular Matrix Disassembly - GO:0022617; Autophagy - hsa04140.

**Tracking of cell migration on nanopatterned substrates**

The nanopatterned substrate, consisting of parallel nanoridges that are 800 nm wide, 800 nm high, and spaced 800 nm apart, was fabricated on top of a glass coverslip as previously described[5]. The coverslip was then mounted at the bottom of an 8-well cell culture chamber (MATTEK). Subsequently, the nanopatterned 8-well chamber was coated with 5 μg/ml laminin at 37 ºC for 2 hours prior to cell seeding. Cells were plated at a low density to facilitate isolated movements, which were observed by time-lapse microscopy. Phase-contrast images were automatically recorded using a 10× objective with SlideBook 6 software for 6 hours at 10-minute intervals. Cells were manually tracked using ImageJ. Migration data were analyzed using a previously published VB program[6]. The number of cells tracked for each patient's primary cells are as follows: Patient 551 - 113 cells; Patient 580 - 120 cells; Patient 601 - 94 cells; Patient 559 - 200 cells; Patient 483 - 130 cells.

**Quantification and statistical analysis**

All statistical tests were performed with Prism (GraphPad) and are described as follows. For all figures, $*p < 0.05$, $**p < 0.01$, $***p < 0.001$, $****p < 0.0001$. he same notations are applied to adjusted p-values. For all data, replicates are shown as individual points, and data shown in the manuscript are three biological replicates with the exception of scRNA-seq, multiplex analysis of cytokines, tracking tumoroid size growth, and IHC quantification of GBM tissues from human patients. Comparisons of two conditions were evaluated using the Two-Tailed t-test or multiple t-tests with two-stage linear step-up procedure of Benjamini, Krieger and Yekutieli. All comparisons involving more than two groups were performed using a One-Way ANOVA with Tukey multiple comparison test.

1. Bai, H., Harmanci, A.S., Erson-Omay, E.Z., Li, J., Coskun, S., Simon, M., Krischek, B., Ozduman, K., Omay, S.B., Sorensen, E.A., et al. (2016). Integrated genomic characterization of IDH1-mutant glioma malignant progression. Nat Genet *48*, 59-66. 10.1038/ng.3457.
2. Mishra-Gorur, K., Barak, T., Kaulen, L.D., Henegariu, O., Jin, S.C., Aguilera, S.M., Yalbir, E., Goles, G., Nishimura, S., Miyagishima, D., et al. (2023). Pleiotropic role of TRAF7 in skull-base meningiomas and congenital heart disease. Proc Natl Acad Sci U S A *120*, e2214997120. 10.1073/pnas.2214997120.
3. Levchenko, A., Ewald, A.J., Lee, S.H., Ellison, D., and Kang, T.-Y. (2019). 3D Analysis of Multi-cellular Responses to Chemoattractant Gradients. Journal of Visualized Experiments. 10.3791/59226-v.
4. Kim, S., Lee, H., Chung, M., and Jeon, N.L. (2013). Engineering of functional, perfusable 3D microvascular networks on a chip. Lab on a Chip *13*. 10.1039/c3lc41320a.
5. Kim, D.H., Han, K., Gupta, K., Kwon, K.W., Suh, K.Y., and Levchenko, A. (2009). Mechanosensitivity of fibroblast cell shape and movement to anisotropic substratum topography gradients. Biomaterials *30*, 5433-5444. 10.1016/j.biomaterials.2009.06.042.
6. Gorelik, R., and Gautreau, A. (2014). Quantitative and unbiased analysis of directional persistence in cell migration. Nat Protoc *9*, 1931-1943. 10.1038/nprot.2014.131.
